# A new GRAB sensor reveals differences in the dynamics and molecular regulation between neuropeptide and neurotransmitter release

**DOI:** 10.1101/2024.05.22.595424

**Authors:** Xiju Xia, Yulong Li

## Abstract

The co-existence and co-transmission of neuropeptides and small molecule neurotransmitters in the same neuron is a fundamental aspect of almost all neurons across various species. However, the differences regarding their *in vivo* spatiotemporal dynamics and underlying molecular regulation remain poorly understood. Here, we developed a GPCR-activation-based (GRAB) sensor for detecting short neuropeptide F (sNPF) with high sensitivity and spatiotemporal resolution. Furthermore, we explore the differences of *in vivo* dynamics and molecular regulation between sNPF and acetylcholine (ACh) from the same neurons. Interestingly, the release of sNPF and ACh shows different spatiotemporal dynamics. Notably, we found that distinct synaptotagmins (Syt) are involved in these two processes, as Syt7 and Sytα for sNPF release, while Syt1 for ACh release. Thus, this new GRAB sensor provides a powerful tool for studying neuropeptide release and providing new insights into the distinct release dynamics and molecular regulation between neuropeptides and small molecule neurotransmitters.

## INTRODUCTION

Neurons typically utilize two primary classes of signaling molecules for transmitting information: neuropeptides like oxytocin (OT), somatostatin (SST), and corticotropin-releasing factor (CRF), alongside small molecule neurotransmitters such as acetylcholine (ACh), glutamate (Glu), and γ-aminobutyric acid (GABA)^1^. Neuropeptides and small molecule neurotransmitters are typically stored in large dense-core vesicles (LDCVs) and synaptic vesicles (SVs)^2^, respectively, which likely have distinct properties that govern their activity-dependent release^3–5^. Moreover, the classic study demonstrated that the neuropeptide and the small molecule neurotransmitter induced slow and fast excitatory postsynaptic potential respectively in sympathetic ganglia^6^. Interestingly, the presence of both neuropeptides and small molecule neurotransmitters in the same neuron is common in almost all neurons to a wide range of species^3,4,7,8^, providing a diverse set of modulatory mechanisms that can operate on distinct spatial and/or temporal scales, thereby enabling complex behaviors such as the flight response, sleep, learning, and social behaviors^5,6,9–13^. However, most of the previous studies examining the release of neuropeptides and the release of small molecule neurotransmitters were conducted separately in distinct cell types; therefore, the potential similarities and/or differences in their spatiotemporal dynamics and their underlying molecular regulation within the same neuron have remained poorly understood.

*Drosophila* is an excellent model organism for studying the regulation of neuropeptides and small molecule neurotransmitters *in vivo* due to its less redundant genome compared to mammals, as well as its well-developed genetic tools and database^14,15^. Short neuropeptide F (sNPF), an ortholog of neuropeptide Y (NPY) in vertebrates, is one of important neuropeptides in *Drosophila*, which is critical for feeding, metabolism, sleep and glucose homeostasis^16–21^. Notably, transcriptomics data revealed the presence of both the neuropeptide sNPF and the small molecule neurotransmitter ACh in Kenyon cells (KCs) in the *Drosophila* mushroom body (MB)^22,23^. These cells function as the olfactory learning center, and both sNPF and ACh have been shown to be important for learning and memory^23–25^. Thus, KCs provide an ideal platform for studying the “co-transmission” of neuropeptide and small molecular neurotransmitter within the same neuron. Previously, we have developed, characterized, and utilized a G protein-coupled receptor (GPCR) activation‒based (GRAB) ACh sensor (GRAB_ACh3.0_) for use in *Drosophila* studies *in vivo*^26,27^; however, a suitable tool for detecting sNPF release *in vivo* is currently unavailable.

Several methods have been developed for detecting neuropeptide release *in vivo*, each with its own advantages and disadvantages. Microdialysis has been widely used to measure the dynamics of neuropeptide release in the mammalian brain^28^; however, this technique is invasive and has low spatiotemporal resolution due to the relatively large embedded probe (∼200 µm diameter) and low sampling rate (requiring 5–10 minutes per sample). Alternatively, neuropeptides tagged with either a fluorescent protein or fluorogen-activating protein (FAP) have been used to track the release of neuropeptides or to monitor the fusion of LDCVs; examples include GFP-tagged rat atrial natriuretic peptide (ANP^GFP^)^29^, pHluorin-tagged neuropeptide Y (NPY-pHluorin)^30^, the GCaMP6s-tagged rat atrial natriuretic peptide neuropeptide release reporter (NPRR^ANP^)^31^, and FAP-tagged *Drosophila* insulin-like peptide 2 (Dilp2-FAP)^32^, these reporters offer good cell specificity and sensitivity for neuropeptide detection *in vivo*. However, because the fluorescent tag is usually ∼10–100 times larger than the neuropeptide itself in terms of molecular weight, these reporters do not necessarily reflect the true dynamics of endogenous neuropeptides. Another approach is to fuse the fluorescent tag to the luminal side of an LDCV-specific membrane protein such as cytochrome b561, providing a versatile tool for monitoring neuropeptide release; however, this approach lacks neuropeptide specificity^33^. The Tango GPCR assay can also be used to detect neuropeptide release *in vivo*, but requires a relatively long time for reporter expression and is irreversible^34–36^. Finally, CNiFER (cell-based neurotransmitter fluorescent engineered reporter) biosensors require the implantation of genetically modified cells, making it highly invasive and lacking cell type specificity^37–41^.

Recently, taking advantage of the GRAB strategy, our group and others independently developed several series of genetically encoded fluorescent sensors for detecting small molecule neurotransmitters and mammalian neuropeptides with high specificity and spatiotemporal resolution^26,42–57^. Capitalizing on the scalability of this approach, we therefore developed a GRAB sensor for detecting the *in vivo* dynamics of sNPF in *Drosophila*. By expressing both the sNPF and ACh sensors in KCs in the *Drosophila* MB and performing *in vivo* two-photon imaging, we then measured the spatiotemporal dynamics of both sNPF and ACh release in real time. We found that sNPF release shows distinct spatiotemporal dynamics with ACh release, while both sNPF and ACh release require neuronal synaptobrevin (nSyb). To further investigate the molecular regulation of sNPF and ACh release, we performed CRISPR/Cas9-based screening of the synaptotagmin family of proteins in the KCs and found that sNPF release is largely mediated by Syt7 and Sytα, while ACh release is mainly mediated by Syt1.

## RESULTS

### Development and characterization of GRAB_sNPF_ sensors

To generate a GRAB sensor for detecting sNPF (GRAB_sNPF_), we first replaced the third intracellular loop (ICL3) in the sNPF receptor (sNPFR) with the ICL3-circularly permutated EGFP (cpEGFP) module from the well characterized norepinephrine sensor GRAB_NE1m_^45^ (Fig. 1A). Because the sNPF peptide sequence is highly conserved among Diptera, including flies and mosquitoes^18^ (Fig. S1A), we screened a series of sNPFRs cloned from these genera^58,59^ (Fig. S1B). We then expressed candidate sensors in HEK293T cells and examined their maximum brightness and change in fluorescence (ΔF/F_0_) in response to application of 1 µM sNPF (unless indicated otherwise, we used the *Drosophila* sNPF2 neuropeptide). The most promising candidate was based on the *Culex quinquefasciatus* sNPFR, which has the highest response and relatively high brightness. We named this sensor sNPF0.1 and utilized it for further optimization (Fig. 1B and Fig. S1A, B). After optimizing the replacement sites, performing site-directed mutagenesis on cpEGFP and linker sequences between cpEGFP and the GPCR, we obtained GRAB_sNPF1.0_ (hereafter referred to as sNPF1.0), which has a peak ΔF/F_0_ of ∼350% in response to sNPF application (Fig. 1C and Fig. S1C, D). Structural data suggested that D287^6.59^ serves as a predicted binding site between NPY, a vertebrate ortholog of sNPF, and its receptor Y_1_R^58,60^. Based on this, we developed an sNPF-insensitive sensor, sNPFmut, by introducing the arginine mutagenesis in the corresponding site D302^6.59^ of sNPF1.0 (Fig. 1C and Fig. S1D). When expressed in HEK293T cells, sNPF1.0 traffics to the plasma membrane (Fig. 1D) and has a concentration-dependent increase in fluorescence in response to sNPF, with an EC_50_ of 64 nM (Fig. 1E); in contrast, sNPFmut showed non-detectable response to sNPF at all concentrations tested (Fig. 1E).

**Fig. 1.**
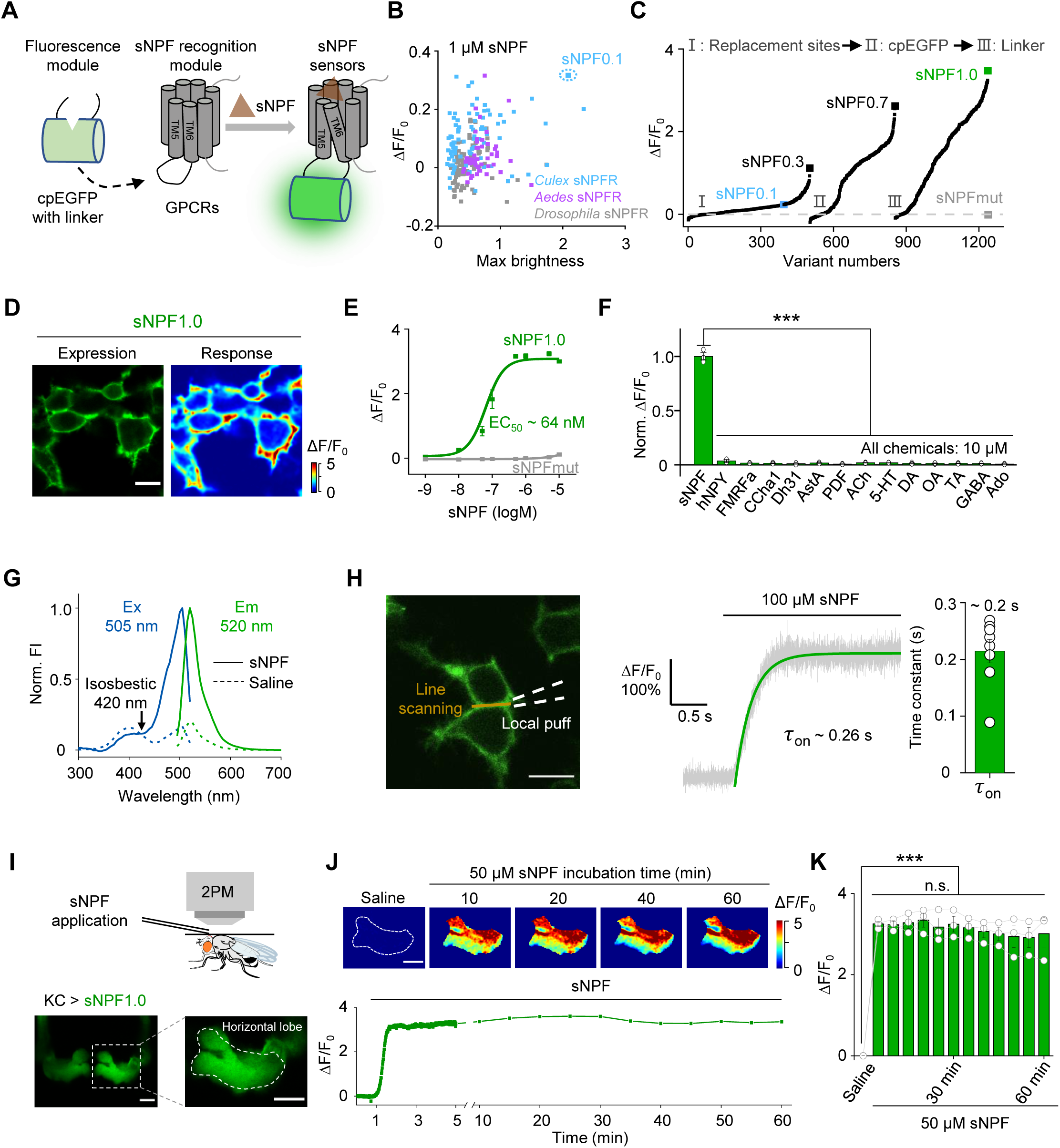
Development and *in vitro* and *in vivo* characterization of GRAB_sNPF_ sensors. (A) Schematic diagram depicting the principle behind the GRAB_sNPF_ sensors in which ICL3 in the sNPF receptor (sNPFR) is replaced with cpEGFP and linker from GRAB_NE1m_. Binding of sNPF to the sensor induces a conformational change that increases the fluorescence signal. (B) Selection of a candidate sensor for further optimization in HEK293T cells by screening sNPFRs cloned from the indicated species. The candidate sensor with the strongest response to 1 μM sNPF, GRAB_sNPF0.1_ (sNPF0.1), is indicated. (C) Optimization of the replacement site, key amino acids in cpEGFP, and linkers between cpEGFP and GPCR in GRAB_sNPF_ sensors based on sNPF0.1, yielding increasingly more responsive sensors. The sensor with the strongest response to 1 μM sNPF, GRAB_sNPF1.0_ (sNPF1.0), is indicated. (D) Representative fluorescence image of sNPF1.0 (left) and pseudocolor image (right) showing the change in sNPF1.0 fluorescence in HEK293T cells expressing sNPF1.0 in response to 1 μM sNPF. Scale bar, 10 μm. (E) Dose–response curves measured in HEK293T cells expressing sNPF1.0 or sNPFmut, with the corresponding EC_50_ values; n = 3 wells with 200-400 cells per well. (F) Summary of normalized ΔF/F_0_ measured in sNPF1.0-expressing HEK293T cells in response to the indicated compounds; n = 3 wells with 200-400 cells per well. sNPF, *Drosophila* short neuropeptide F; hNPY, human neuropeptide Y; FMRFa, FMRFamide; CCHa1, CCHamide 1; Dh31, diuretic hormone 31; AstA, allatostatin A; PDF, pigment-dispersing factor; ACh, acetylcholine; 5-HT, 5-hydroxytryptamine; DA, dopamine; OA, octopamine; TA, tyramine; GABA, gamma-aminobutyric acid; Ado, adenosine. (G) One-photon excitation (ex) and emission (em) spectra of sNPF1.0 measured in the absence and presence of sNPF. The isosbestic point and excitation and emission peaks are indicated. FI, fluorescence intensity. (H) Summary of the kinetics of the sNPF1.0 response. Left: illustration of the local puffing system. Middle: a representative response trace. Right: group data summarizing τ_on_; n = 8 cells from 3 cultures. Scale bar, 10 μm. (I) Schematic illustration (top) and fluorescence images (bottom) of a transgenic fly expressing sNPF1.0 in MB KCs. Scale bar, 25 μm. (J) Representative pseudocolor images (top) and trace (bottom) of ΔF/F0 in response to a 1-hour perfusion of 50 μM sNPF in a transgenic fly expressing sNPF1.0 in MB KCs. Scale bar, 25 μm. (K) Summary of ΔF/F0 measured in response to 50 μM sNPF at the indicated times; n = 3 flies. Data are shown as mean ± s.e.m. in h, with the error bars or shaded regions indicating the s.e.m. \*\*\**P* < 0.001, \*\**P* < 0.01, \**P* < 0.05, and n.s., not significant.

We then characterized the specificity, spectral properties, and kinetics of sNPF1.0 expressed in HEK293T cells. sNPF1.0 has high specificity for sNPF, with virtually no response elicited by a wide range of neuropeptides and small molecule neurotransmitters (Fig. 1F). Moreover, sNPF1.0 can detect other sNPF analogs and homologs from *Drosophila* and *Culex*, with similar peak responses but with EC_50_ values ranging from 23 nM to 1.7 μM (Fig. S2A-C). We measured one-photon spectral properties of sNPF1.0, with peak excitation and emission wavelengths of 505 nm and 520 nm, respectively (Fig. 1G), as well as a two-photon excitation peak at 930 nm (Fig. S2D). With respect to the sensor’s activation kinetics, we measured an average rise time constant (τ_on_) of approximately 0.2 s (Fig. 1H). Finally, we confirmed that sNPF1.0 shows no detectable downstream coupling by measuring G protein‒dependent pathways and β-arrestin recruitment, although wild-type *Culex* sNPFR activated both signaling pathways in response to sNPF (Fig. S2E, F).

Next, we evaluated the ability of sNPF1.0 to detect sNPF *in vivo* by expressing sNPF1.0 in KCs in the *Drosophila* MB. Using two-photon imaging, we then measured the change in sNPF1.0 fluorescence in response to sNPF application (Fig. 1I). Application of 50 μM sNPF induced a robust increase in sNPF1.0 fluorescence that was stable for at least 60 min (Fig. 1J, K), suggesting minimal internalization or desensitization of the sensor *in vivo*, and showing that sNPF1.0 is suitable for long-term imaging.

### GRAB_sNPF_ reports endogenous sNPF release *in vivo*

Then, we examined whether sNPF1.0 can detect the release of endogenous sNPF. We expressed sNPF1.0 pan-neuronally under the control of nSyb-Gal4, and mainly focused on the fluorescent change in MB, due to previous studies showed that sNPF is highly expressed in KCs in the *Drosophila* MB^19,61^. We found that high K^+^ induced an increase in sNPF1.0 fluorescence in the horizontal lobe of MB (Fig. 2A-C). In contrast, no apparent response to high K^+^ was measured in sNPF1.0-expressing sNPF-knockout (sNPF-KO) flies. However, the exogenous application of sNPF still elicited a robust response in these flies, indicating the sensor expression was unaffected (Fig. 2B-C).

**Fig. 2.**
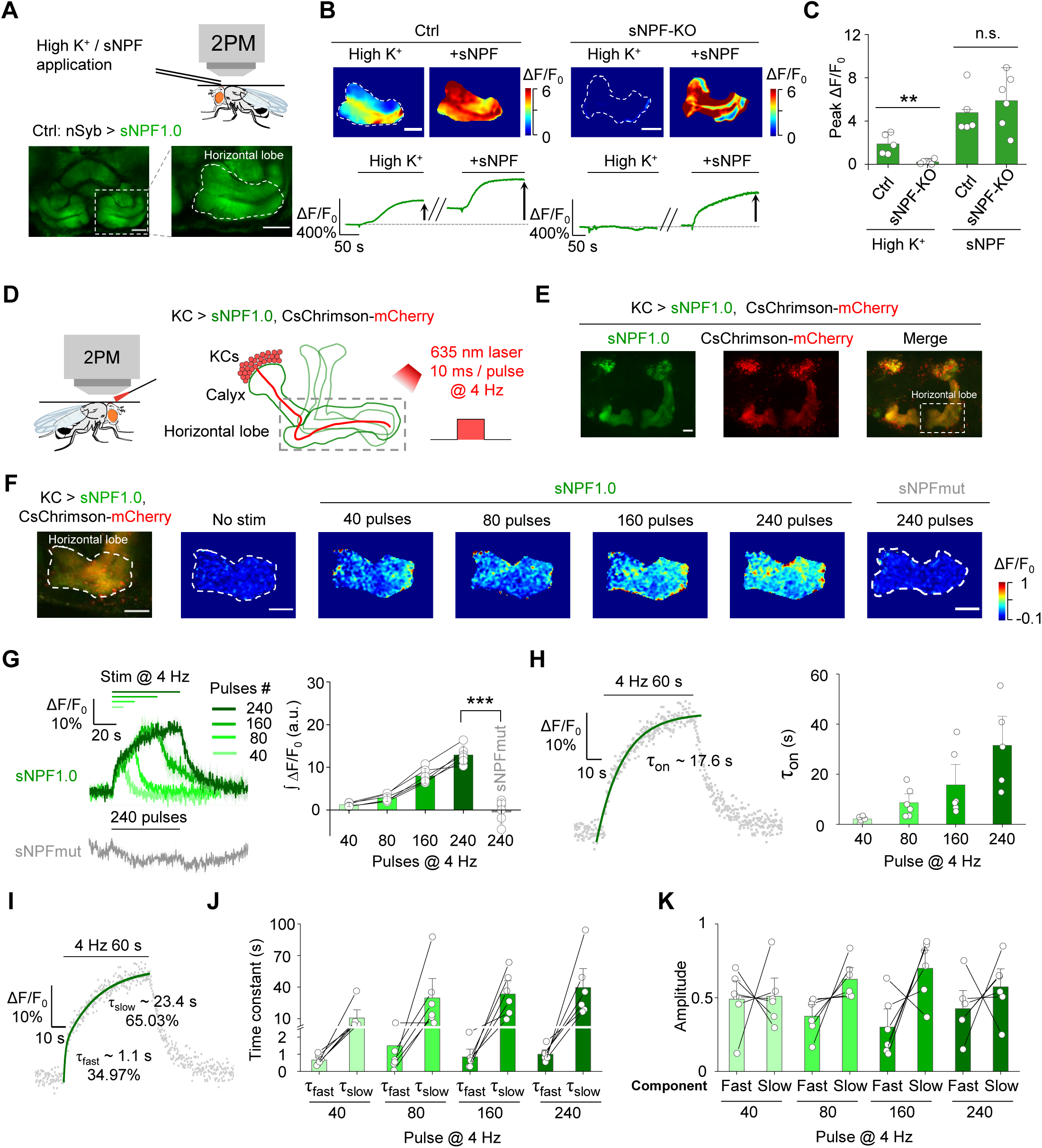
The sNPF1.0 GRAB sensor can detect sNPF release *in vivo*. (A) Schematic diagram (top) and representative fluorescence images (bottom) of sNPF1.0 expressed in the horizontal lobe in the *Drosophila* MB. Scale bar, 25 μm. (B) Representative pseudocolor images (top) and traces (bottom) of sNPF1.0 expressed in control flies (left) and sNPF KO flies (right); where indicated, high K^+^ and sNPF were applied. Scale bars, 25 μm. (C) Summary of peak ΔF/F_0_ measured in the indicated flies in response to high K^+^ and sNPF; n = 5-6 flies each. (D) Schematic illustration depicting the experimental setup. CsChrimson-mCherry and sNPF1.0 were expressed in KCs in the *Drosophila* MB, and 635-nm laser light pulses were used to optogenetically activate the KCs. (E) Representative fluorescence images of sNPF1.0 and CsChrimson-mCherry in the MB; the horizontal lobe is indicated by the dashed white box. Scale bar, 25 μm. (F) Fluorescence image of sNPF1.0 and CsChrimson-mCherry in the horizontal lobe in KCs (left-most image) and representative pseudocolor images (right) of the fluorescence responses of sNPF1.0 and sNPFmut to the indicated number of 635-nm laser pulses applied at 4 Hz. Scale bars, 25 μm. (G) Traces (left) and summary (right) of the fluorescence responses of sNPF1.0 and sNPFmut; n = 6 flies each. (H) sNPF1.0 fluorescence was measured before, during, and after a 240-pulse train of 635-nm light. The rise phase was fitted with a single-exponential function (left), and the time constants (τ_on_) are summarized on the right; n = 6 flies. (I) sNPF1.0 fluorescence was measured before, during, and after a 240-pulse train of 635-nm light, and the rise phase was fitted with a double-exponential function. (J and K) Summary of the fast and slow time constants (J) and relative amplitudes (K) measured as shown in (I); n = 6 flies Data are shown as mean ± s.e.m. in g, with the error bars or shaded regions indicating the s.e.m. \*\**P* < 0.01, \**P* < 0.05, and n.s., not significant.

To achieve cell autonomous and high temporal control of endogenous sNPF release in KCs, we utilized CsChrimson to activate KCs, and measured sNPF release in response to optogenetic activation^62^ in the axonal region (i.e., the horizontal lobe) (Fig. 2D, E) and the dendritic region (i.e., the calyx) (Fig. S3A) of KCs *in vivo*. We found that optogenetic stimulation evoked time-locked and pulse number‒dependent sNPF release in both regions (Fig. 2F-H and Fig. S3A-C). In contrast, no detectable response was observed in sNPF-mut expressed flies (Fig. 2G). The rise time constant (τ_on_) in the axonal and dendritic regions ranged from 2.1–26.9 s and 4.3–19.9 s, respectively, with time constants correlated with increasing pulse numbers in both regions (Fig. 2H and Fig. S3D). Interestingly, the rising phase of the sNPF1.0 signal was best fit with a double-exponential function, reflecting the existence of both a fast rising phase and a relatively slow rising phase (Fig. 2I-K and Fig. S3E-G).

Taken together, these results indicate that sNPF1.0 is able to report the endogenous sNPF release specifically and is suitable to study the spatiotemporal dynamics of sNPF release *in vivo*.

### GRAB sensors reveal spatially distinct patterns of sNPF and ACh release from KCs

Many neurons—including KCs in the *Drosophila* MB—produce and release both neuropeptides and small molecule neurotransmitters. To compare their spatiotemporal dynamics, we therefore measured the release patterns of sNPF and ACh by optogenetically activating KCs in MB. Specifically, we expressed either sNPF1.0 or the ACh sensor ACh3.0^26^ along with CsChrimson in KCs (Fig. 3A). To avoid potential interference induced by activating other neurons through ACh release, we included the nicotinic ACh receptor blocker mecamylamine (Meca) throughout these experiments. We found that optogenetic stimulation of KCs induced sNPF release in the axons (horizontal lobe), dendrites (calyx), and soma regions; in contrast, ACh release was restricted to the axonal and dendritic regions (Fig. 3B-E and Fig. S4). In addition, the levels of both sNPF release and ACh release from the axons were significantly higher compared to their release from the dendrites (Fig. 3E). These results indicate that sNPF and ACh have different spatial release patterns from KCs.

**Fig. 3.**
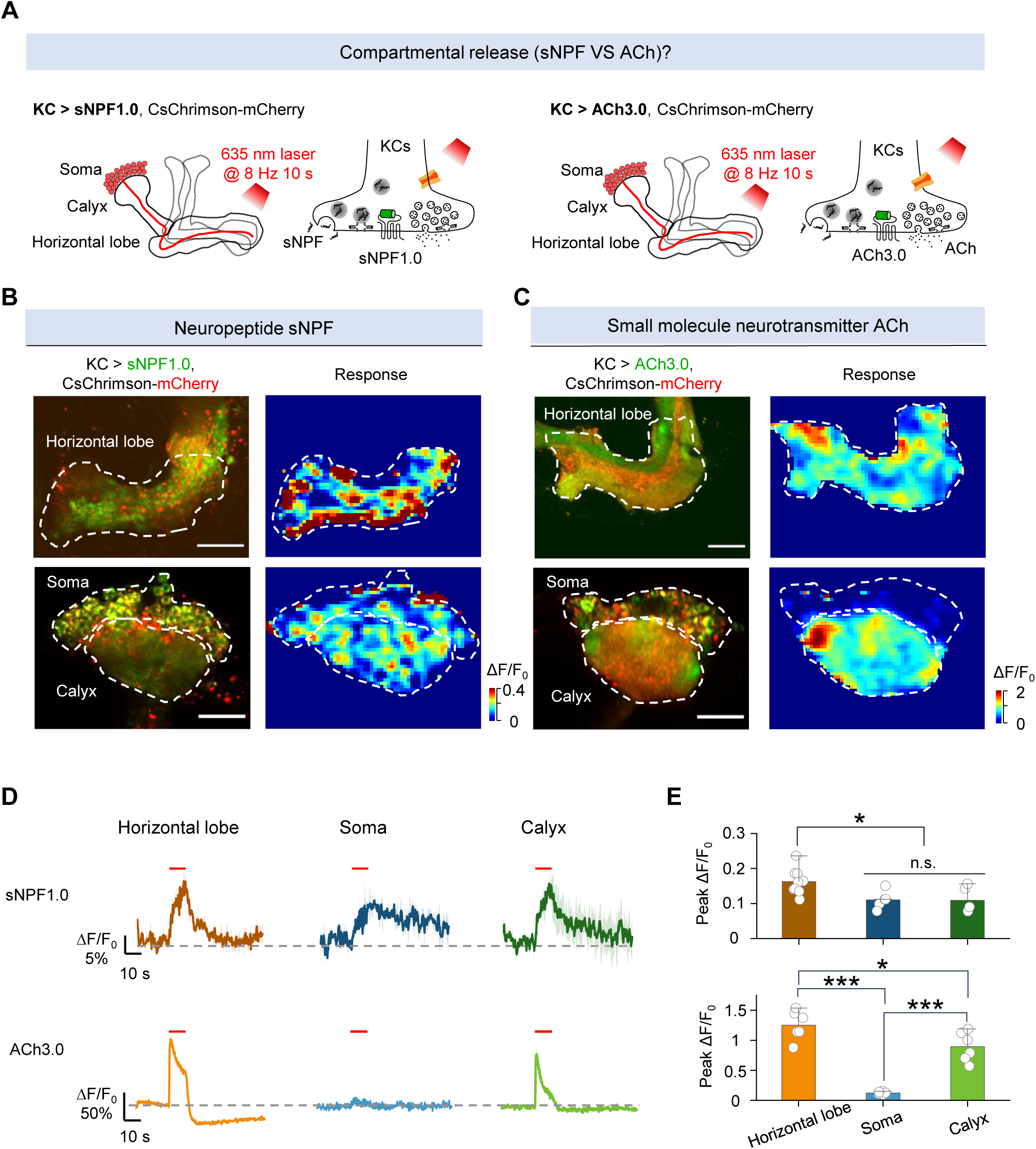
The sNPF1.0 and ACh3.0 GRAB sensors reveal spatial difference in release between sNPF and ACh. (A) Schematic diagram depicting the KC regions in the *Drosophila* MB, which can be divided into the axon (horizontal lobe), dendrite (calyx), and soma regions. Also shown is the strategy for imaging sNPF and ACh release in the MB using sNPF1.0 and ACh3.0, respectively. The 100 μM nAChR antagonist mecamylamine (Meca) was present throughout these experiments. (B and C) Representative fluorescence images (left columns) and pseudocolor images (right columns) showing the change in sNPF1.0 (B) and ACh3.0 (C) fluorescence in response to 80 light pulses delivered at 8 Hz. The top rows show the horizontal lobe, and the bottom rows show the calyx and soma regions (dashed outlines). Scale bars, 25 μm. (D and E) Representative traces (D) and quantification (E) of the change in sNPF1.0 (D, top) and ACh3.0 (D, bottom) fluorescence in response to 80 light pulses delivered at 8 Hz. Data are shown as mean ± s.e.m. in d, with the error bars or shaded regions indicating the s.e.m. \*\*\**P* < 0.001, \**P* < 0.05, and n.s., not significant.

### GRAB sensors reveal distinct activity-dependent dynamics underlying sNPF and ACh release

Having shown the differences in the spatial release patterns between sNPF and ACh, we next asked whether differences exist in release probability and the temporal dynamics of their release. Although it is generally believed that neuropeptide release is slower compared to the release of small molecule neurotransmitter^63^, this has not been examined directly by measuring the release of these two types of signaling molecules within the same cell type *in vivo*. Given that axons exhibited a higher release probability compared to other neuronal compartments (Fig. 3E), we examined the kinetics and temporal profiles of sNPF and ACh release in the horizontal lobe in flies expressing CsChrimson together with either sNPF1.0 or ACh3.0 (Fig. 4A). We found that light pulses generated an sNPF1.0 signal that had slower rise and decay kinetics (τ_on_: 0.94–4.4 s; τ_off_: 4.9–7.2 s) compared to the ACh3.0 signal (τ_on_: 0.13–0.24 s; τ_off_: 1.1–1.4 s) (Fig. 4B-G). Given that the activation kinetics of sNPF1.0 and ACh3.0 sensors are ∼0.2 s (Fig. 1H) and ∼0.15 s^26^, respectively, when compared to the ACh signal, the physiologically slower kinetics of the sNPF signal induced by optogenetic stimulations suggest a distinction between the release of neuropeptides and small molecule neurotransmitters from the same neurons.

**Fig. 4.**
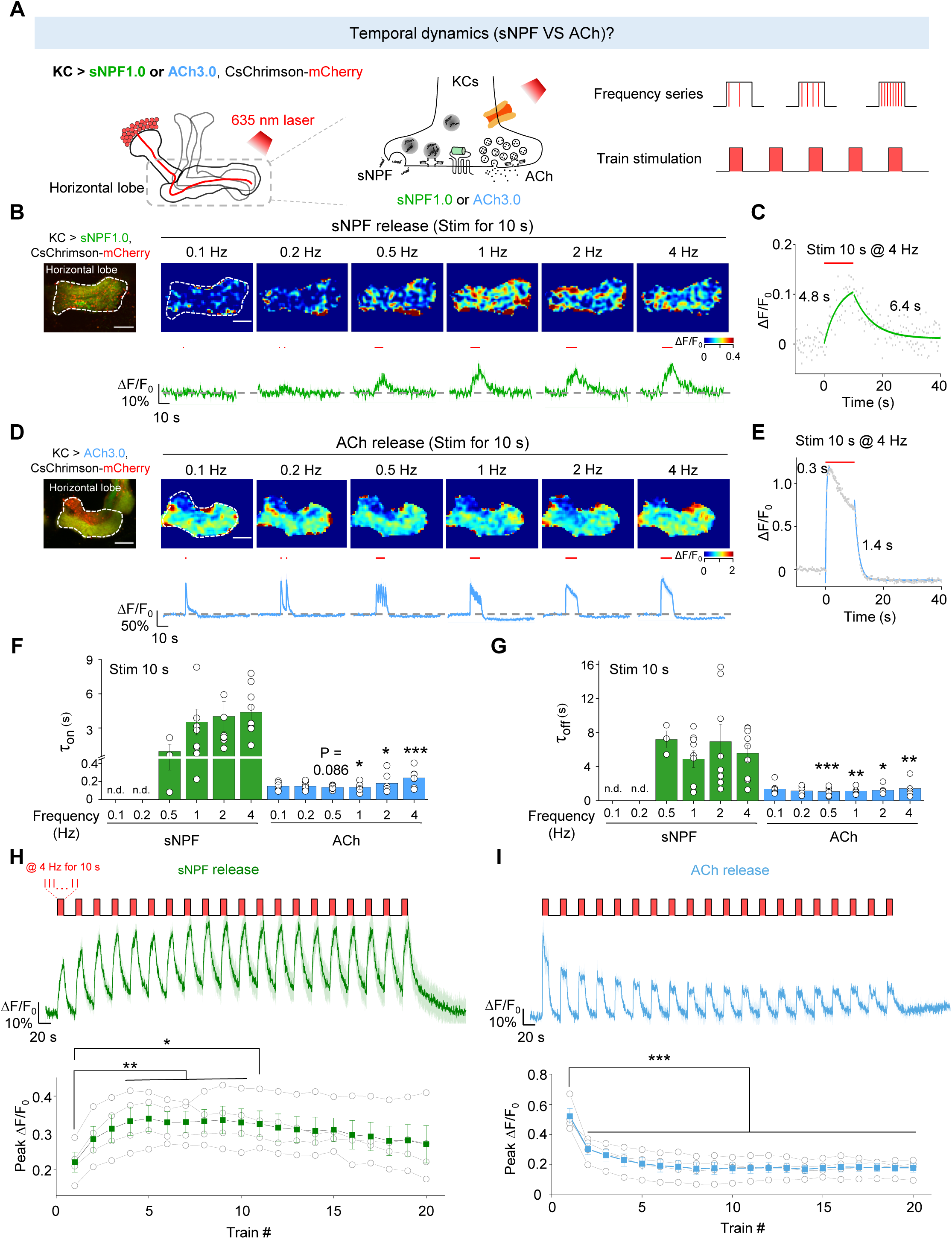
The sNPF1.0 and ACh3.0 GRAB sensors reveal distinct activity-dependent properties for sNPF and ACh release. (A) Schematic diagram depicting the strategy for measuring the temporal dynamics of sNPF or ACh release in the horizontal lobe using sNPF1.0 and ACh3.0, respectively. The 100 μM nAChR antagonist mecamylamine (Meca) was present throughout these experiments. (B and D) Representative fluorescence image (top left), pseudocolor images (top right), and traces (bottom right) of the change in sNPF1.0 (B) and ACh3.0 (D) fluorescence in response to the indicated light stimuli (red bars). Scale bars, 25 μm. (C and E) Example traces showing the change in sNPF1.0 (C) and ACh3.0 (E) fluorescence before, during, and after the indicated light stimuli; the rise and decay phases are each fitted with a single-exponential function. (F and G) Summary of the rise (F) and decay (G) time constants (τ_on_ and τ_off_) measured for the change in sNPF1.0 and ACh3.0 fluorescence in response to the indicated light stimuli. (H and I) Individual traces (top) and summary (bottom) of the change in sNPF1.0 (H) and ACh (I) fluorescence in response to the indicated light stimuli. Data are shown as mean ± s.e.m. in B, D, H and I, with the error bars or shaded regions indicating the s.e.m. \*\*\**P* < 0.001, \*\**P* < 0.01, and \**P* < 0.05.

Moreover, the sNPF-containing LDCVs have a high release threshold since the peak sNPF1.0 signal showed the light pulse frequency-dependent manner (Fig. 4B), whereas the peak ACh3.0 signal was largely unaffected by stimulation frequency (Fig. 4D), suggesting a large difference in the initial release probability.

When multiple stimuli were delivered within a short interval, the release of neurotransmitter or neuromodulator can be either enhanced or depressed relative to that induced by the initial stimulus^64^. This phenomenon is named as short-term plasticity, which is implicated in various physiological functions and pathological conditions, such as learning, memory and some psychiatric disorders^64,65^. To further test the short-term plasticity, we examined the release pattern of sNPF and ACh and found that applying more light pulses at a fixed frequency (1 Hz) potentiated the sNPF1.0 signal, but depressed the ACh3.0 signal (Fig. S5), suggesting post-tetanic potentiation of neuropeptide release. What’s more, when we applied a stimulation protocol consisting of repeated trains of light pulses, the results showed that sNPF release was potentiated during this stimulation protocol (Fig. 4H), while ACh release was attenuated (Fig. 4I).

Taken together, the above results suggest that sNPF-containing LDCVs have a low release probability, and ACh-containing SVs have a high release probability. In addition, sNPF release has slower kinetics compared to ACh release and shows distinct short-term plasticity with ACh release.

### GRAB sensors reveal that sNPF and ACh reside in vesicle pools with distinct properties

Vesicle pools play a critical role in presynaptic physiology, particularly with respect to release probability and determining synaptic strength, the sizes of vesicle pools are dynamically changing in response to stimuli^66^. To evaluate the dynamics of the vesicle pools containing sNPF and ACh in KCs, we used either continuous stimuli or trains of stimuli to activate KCs (Fig. 5A, B); as above, we included Meca throughout these experiments. Firstly, to examine the dynamics of vesicle pools in response to the long continuous stimuli, we applied a 40-pulse train, followed by a 30-min train of 7200 pulses, followed by several brief stimuli applied at an increasing interval (Fig. 5C). We found that the sNPF1.0 signal initially decreased slightly but was relatively stable during the 30-min stimulation period and the subsequent brief stimuli (Fig. 5C, E). In contrast, the ACh3.0 signal decreased rapidly during the 30-min stimulation period, but recovered during the subsequent brief stimuli (Fig. 5D, F). These data suggest that sNPF resides in a large pool of releasable vesicles so that sNPF release can be maintained with a low release probability for a relatively long period; in contrast, ACh resides in a smaller releasable pool that is rapidly released with a high release probability, but can recover relatively quickly.

**Fig. 5.**
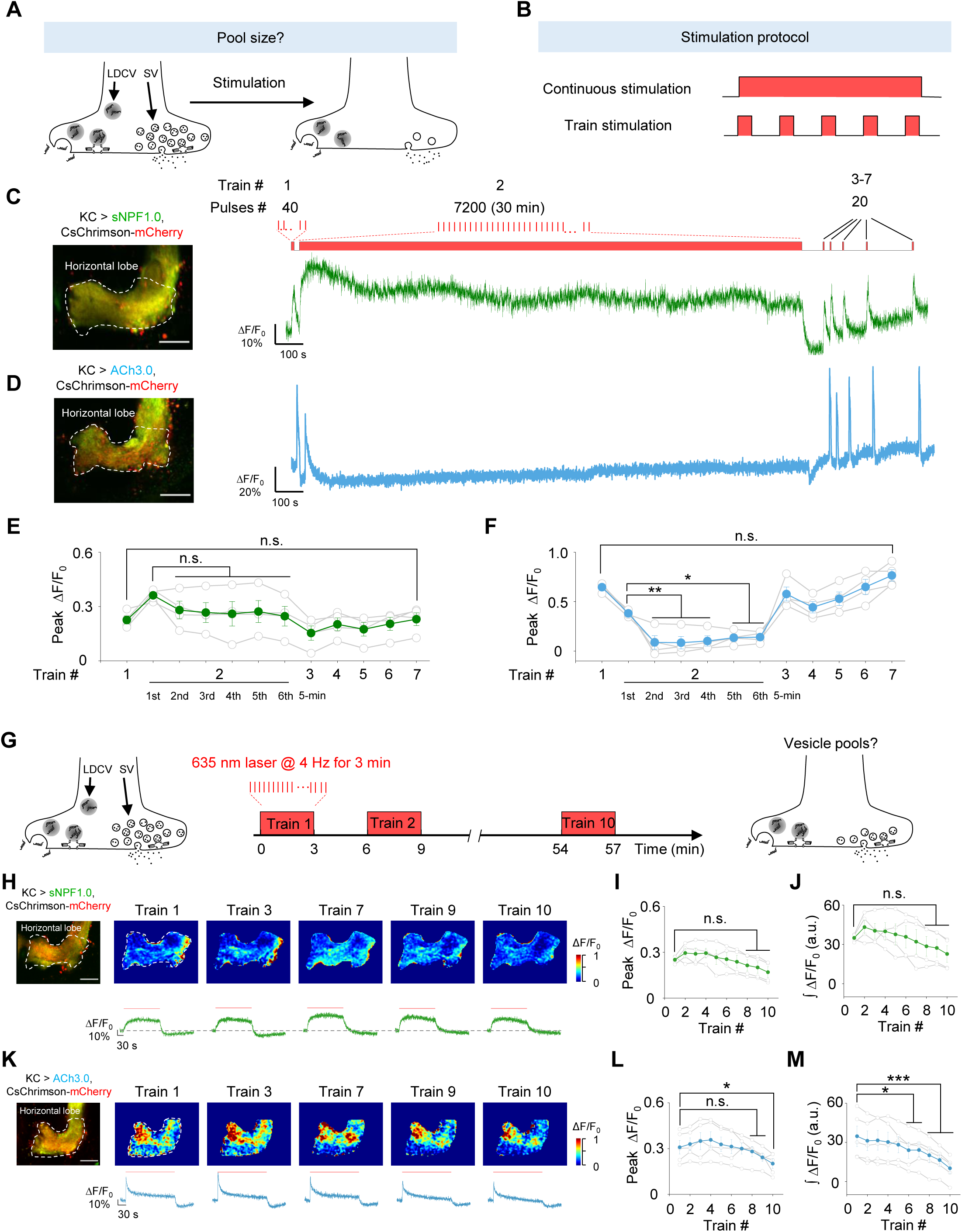
The sNPF1.0 and ACh3.0 GRAB sensors reveal distinct pools of sNPF- and ACh-containing vesicles. (A and B) Schematic diagram depicting the experimental strategy (A) and stimulation protocol (B) used to study the size of vesicle pools containing sNPF and ACh. The 100 μM nAChR antagonist mecamylamine (Meca) was present throughout these experiments. (C and D) Representative fluorescence image (left) and traces (right) of the change in sNPF1.0 (C) and ACh3.0 (D) fluorescence in response to the indicated light stimuli. Scale bars, 25 μm. (E and F) Summary of peak ΔF/F0 measured for sNPF1.0 (E) and ACh3.0 (F); n = 4 flies each. (G) Schematic diagram depicting the strategy for studying vesicle pools containing sNPF and ACh. (H and K) Representative fluorescence images (top left), pseudocolor images (top right), and traces (bottom right) of the change in sNPF1.0 (H) and ACh3.0 (K) fluorescence in response to the indicated trains of light; n = 4 flies. (I and J) Summary of the peak (I) and integrated (J) change in sNPF1.0 (H) fluorescence in response to the indicated trains of light; n = 4 flies. Scale bars, 25 μm. (L and M) Summary of the peak (L) and integrated (M) change in ACh3.0 (K) fluorescence in response to the indicated trains of light; n = 4 flies. Data are shown as mean ± s.e.m. \*\*\**P* < 0.001, \**P* < 0.05, and n.s., not significant.

Next, to further investigate the dynamics of the vesicle pools containing sNPF and ACh during the discontinuous stimuli, we delivered 10 trains of light pulses with a 3-min interval while measuring sNPF or ACh release in the horizontal lobe (Fig. 5G). The results showed a relatively stable peak and integrated response for both sNPF and ACh release in response to these 10 trains (Fig. 5H-M). Such a relative stable response could be attributed to the vesicle pools recovering during each 3-min interval and/or the presence of a relatively large vesicle pool that can maintain release during intense stimulation.

### GRAB sensors reveal that sNPF and ACh release are mediated by overlapping and distinct molecular mechanisms

Both SVs and LDCVs require soluble N-ethylmaleimide-sensitive factor attachment receptor (SNARE) complexes for vesicle fusion^67,68^. In *Drosophila*, neuronal synaptobrevin (nSyb) is a core component of the SNARE complex and is required for the release of small molecule neurotransmitters^69^. In contrast, whether the same SNARE proteins mediate the release of both sNPF and ACh in the same neuron is an open question.

To determine whether nSyb mediates the release of ACh and/or sNPF in KCs, we expressed tetanus toxin light chain (Tetxlc) in KCs to specifically cleave nSyb^70^ and then measured the effect on ACh and sNPF release. We found that expressing Tetxlc significantly reduced both the high K^+^‒induced sNPF1.0 signal (Fig. 6A) and the optogenetically-induced ACh3.0 signal (Fig. 6B), but had no apparent effect on signals induced by direct application of sNPF and ACh, respectively (Fig. 6A, B). Thus, both sNPF release and ACh release require nSyb.

**Fig. 6.**
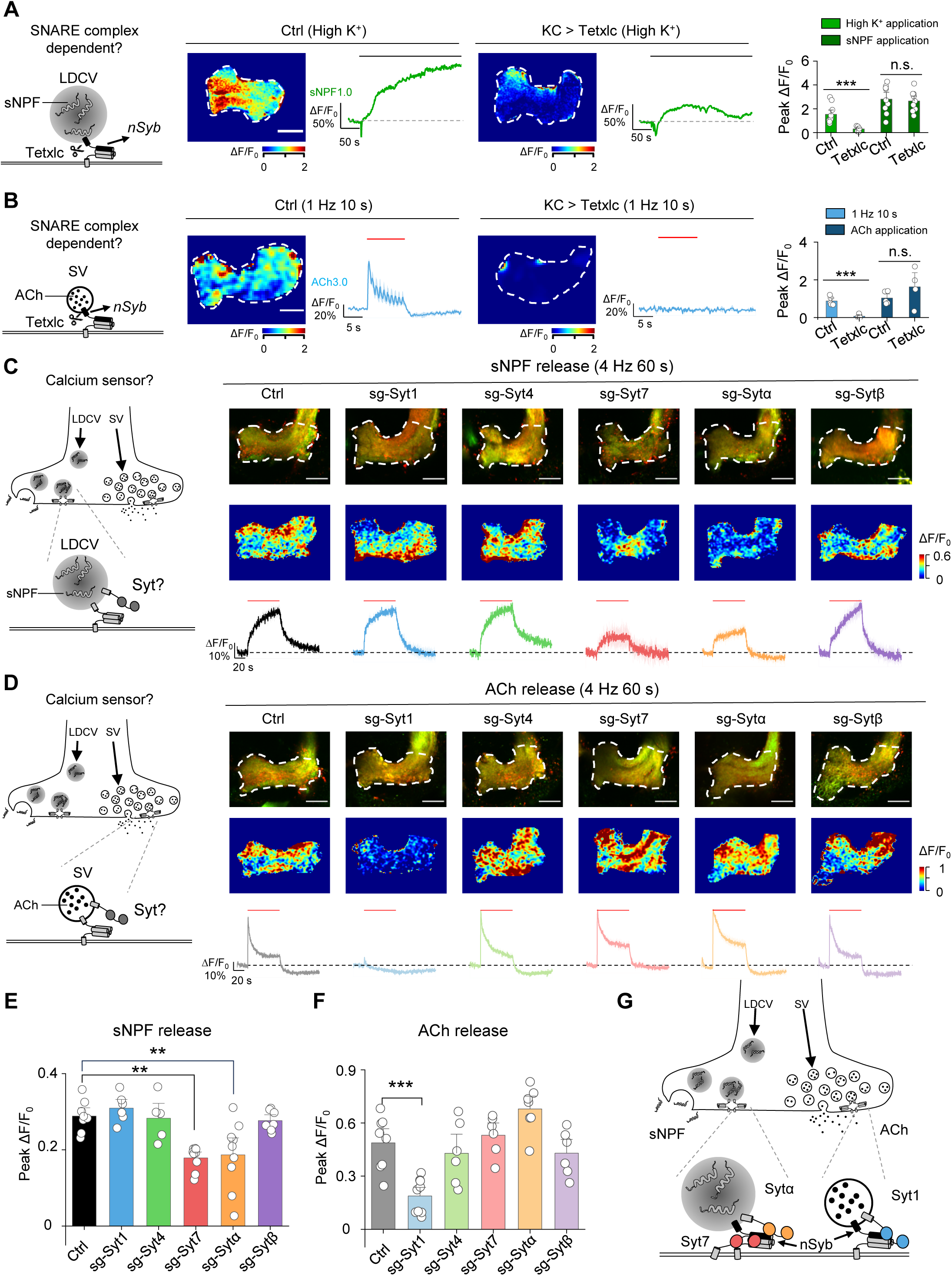
The sNPF1.0 and ACh3.0 GRAB sensors reveal distinct differences in the molecular control of sNPF and ACh release. (A) Left, schematic diagram depicting the release of sNPF via nSyb, a core component of the SNARE complex. Also shown are representative pseudocolor images and traces (middle) and the summary (right) of peak sNPF1.0 ΔF/F0 measured in response to high K^+^ and sNPF application in transgenic flies expressing sNPF1.0 alone (Ctrl) or sNPF1.0 together with the tetanus toxin light chain (Tetxlc) to cleave nSyb. The horizontal lobe is indicated (white dashed outline). Scale bar, 25 μm. (B) Left, schematic diagram depicting the release of ACh via nSyb. Also shown are representative pseudocolor images and traces (middle) and the summary (right) of peak ACh3.0 ΔF/F0 in response to light stimuli and ACh application in transgenic flies expressing ACh3.0 alone (Ctrl) or ACh3.0 together with Tetxlc. Scale bar, 25 μm. (C and D) Left, schematic diagrams depicting the release of sNPF (C) and ACh (D) via synaptotagmins (Syts). Also shown are representative fluorescence images (top right) and pseudocolor images (middle right), and traces (bottom right) of the change in sNPF1.0 (C) and ACh3.0 (D) fluorescence in response to 240 light pulses at 4 Hz in control flies (Ctrl) and in flies in which Syt1, Syt4, Syt7, Sytα, or Sytβ was knocked out using CRISPR/Cas9. (E and F) Summary of the peak change in sNPF1.0 (E) and ACh3.0 (F) fluorescence measured in the indicated flies. Scale bars, 25 μm. (G) Model depicting the shared and distinct proteins that mediate the release of sNPF and ACh in large dense-core vesicles (LDCVs) and synaptic vesicles (SVs), respectively. Data are shown as mean ± s.e.m. in b, c and d, with the error bars or shaded regions indicating the s.e.m. \*\*\**P* < 0.001, \*\**P* < 0.01, and n.s., not significant.

Given that nSyb appears to play a role in the release of both sNPF and ACh, we next investigated the factors that account for the differences in the dynamics of release between sNPF and ACh. The release of neuropeptides and small molecule neurotransmitters (i.e., the fusion of LDCVs and SVs, respectively) is tightly regulated by calcium ions (Ca^2+^)^71^, with synaptotagmins (Syts) serving as the Ca^2+^ sensor, ultimately triggering vesicle fusion^71–73^. With respect to the release of small molecule neurotransmitters in SVs, the function of Syts such as Syt1 and Syt7 has been studied in detail in both vertebrates and invertebrates^74–83^. In contrast, which Syt(s) mediate the release of neuropeptides in LDCVs *in vivo* has not yet been determined.

Syts are a large family of membrane proteins, with seven isoforms present in *Drosophila*. Five of these isoforms— Syt1, Syt4, Syt7, Sytα, and Sytβ—are predicted to bind Ca^2+^ and may therefore regulate the release of neuropeptides and/or small molecule neurotransmitters^84^. To determine which Syt isoform(s) regulate neuropeptide release, we systematically knocked out each of these five Syt isoforms and then measured optogenetically induced sNPF release in KCs using the sNPF1.0 sensor. We utilized a cell type‒specific CRISPR/Cas9-based strategy to knockout each Syt isoform in KCs^85^. Based on this strategy, we generated sgRNA library lines targeting each *Drosophila* Syt isoform, with each isoform targeted by three sgRNAs in one fly line; control flies expressed Cas9 but no sgRNAs. We then performed an imaging screen to compare sNPF release in control flies with that in flies lacking specific Syt isoforms in KCs (Fig. 6C). We found that flies lacking either Syt7 or Sytα had significantly reduced sNPF release in response to optogenetic stimulation (Fig. 6C, E). Surprisingly, knocking out both Syt7 and Sytα did not show a synergistic effect on sNPF release, suggesting that these two Syt isoforms may function in the same pathway (Fig. S6). Finally, we measured ACh release in flies lacking each Syt isoform and found that consistent with the previous studies, knocking out Syt1— but no other isoforms—significantly reduced ACh release (Fig. 6D, F). These results indicate that distinct Syt isoforms regulate different vesicle-release pathways in the same type of neurons, with Syt7 and Sytα mediating neuropeptide release and Syt1 mediating the release of small molecule neurotransmitters (Fig. 6G).

## DISCUSSION

Here, we report the development, characterization, and in *vivo* application of sNPF1.0, a new genetically encoded green fluorescent sensor designed to detect the neuropeptide sNPF. This new sensor has high affinity for sNPF, relatively rapid kinetics, high specificity, and high spatiotemporal resolution. When expressed in *Drosophila*, sNPF1.0 reliably detects the release of sNPF, with a biphasic release pattern during optogenetic stimulation consisting of a fast phase followed by a slow phase. Furthermore, we examined the spatiotemporal patterns of sNPF and ACh release from KCs and found that both sNPF and ACh are released from the axonal and dendritic regions, while sNPF is also released from the soma and has slower kinetics compared to ACh release. Moreover, although both sNPF and ACh require nSyb for their release, our Syt knockout screen revealed that sNPF release is regulated by Sytα and Syt7, whereas ACh release is regulated by Syt1. These differences in Ca^2+^ sensors between sNPF and ACh release may therefore contribute to the observed differences in release kinetics between LDCVs and SVs in the same type of neurons.

### Advantages of GRAB_sNPF_ and its potential applications

Our GRAB_sNPF1.0_ sensor offers several advantages for detecting neuropeptide transmission compared to existing methods. First, this sensor can directly detect the release of endogenous sNPF, making it superior to fluorescent reporter protein‒tagged neuropeptides such as ANP-GFP^29^, NPRR^ANP31^, and Dilp2-FAP^32^. Second, sNPF1.0 has considerably better temporal resolution (τ_on_ ∼0.2 s) compared to microdialysis, which is limited by its relatively slow sampling time (>5 min).

Importantly, sNPF1.0 can be used to measure sNPF release *in vivo* with high specificity, sensitivity, and spatiotemporal resolution. Using sNPF1.0, we explore the dynamics of sNPF release in KCs. In addition to being released from KCs, sNPF can also be released from a wide range of neuron types, playing an important role in regulating various behaviors including circadian rhythms, glucose homeostasis, and body size^16,17,19,20,86^. Moreover, sNPF plays an important role in many insects, including mosquitoes such as *Aedes aegypti*^58^. Therefore, this novel sNPF sensor is suitable for various *in vivo* applications and has potential ability to measure sNPF release in a wide range of behavioral processes and species, providing valuable insights into the regulation of sNPF under a variety of physiological conditions.

### Spatiotemporal dynamics of neuropeptide and small molecule neurotransmitter release from the same type of neurons

The ability of individual neurons to release both neuropeptides and small molecule neurotransmitters is a core feature of neuronal signaling. We found that in contrast to ACh, sNPF can be released from the soma. This was not surprising, given that the somatic release of neuropeptides has been reported in both vertebrates^49,87^ and invertebrates^88^. In *Drosophila*, the somatic release of neuropeptides has been implicated in regulating rhythmic behaviors^88^. This also fits well with structural analyses of neuropeptide release sites in EM sections^89^. Moreover, our results showed that sNPF release kinetics is slower than ACh, which is consistent with the relatively slower fusion of neuropeptide-containing LDCVs compared to neurotransmitter-containing SVs^1^. It also correlated well with the slow and fast excitatory postsynaptic potential induced by the small molecule neurotransmitter and neuropeptide respectively^6^. According to previous literature^90^, different Syt isoforms are known to have different kinetic properties, as Syt1 displayed the fastest disassembly kinetics with Ca^2+^, while Syt7 exhibited the slowest disassembly kinetics. Thus, the observed difference in release kinetics of sNPF and ACh may be attributed to the intrinsic kinetics of distinct Syt. In addition, we found that sNPF release can be maintained for a longer duration than ACh release, suggesting key differences in their respective vesicle pools and indicating that neuropeptides can have broader, longer-lasting effects than small molecule neurotransmitters.

Even after several decades of research, understanding the patterns of neural activity required to drive the release of both neuropeptides and small molecule neurotransmitters from the same neuron remains elusive. Fluorescence sensors can greatly facilitate the analysis of these patterns by detecting the release of neuropeptides and small molecule neurotransmitters under optogenetic-mediated specific activation patterns. Here, we show that sNPF1.0 and ACh3.0 can be used to determine the optogenetic parameters needed to trigger the *in vivo* release of sNPF and ACh, respectively, in the *Drosophila* MB. Notably, trains of optogenetic pulses induced a potentiation of sNPF release, but caused a depression in ACh release, suggesting that distinct processes may underlie the regulation of various phases during complex behaviors. The post-tetanic potentiation of neuropeptide release was also observed in larval *Drosophila* neuromuscular junctions^91^. We also speculate that sNPF-containing LDCVs have a low release probability, and ACh-containing SVs have a high release probability. This finding aligned with the distinct localization patterns of LDCVs and SVs, where SVs tend to cluster near the active zone, while LDCVs are dispersed in remote regions away from the active zone^92^.

### Molecular regulation of neuropeptide release

The Syt family is highly conserved across different species, with *Drosophila* Syt1 and Syt7 being orthologous to the mouse Syt1 and Syt7 genes respectively, furthermore, *Drosophila* Sytα shares the highest similarity to mouse Syt9, Syt10, and Syt3^84^. Despite decades of study, the function of most Syt isoforms with respect to the release of neuropeptides remains poorly understood. To address this question, we systematically screened all five putative Ca^2+^-sensitive Syt isoforms for their role in mediating neuropeptide release in the *Drosophila* MB and found that both Sytα and Syt7 are required for sNPF release. It was correlated well with previous reports, such as Park et al. reported that knocking down Sytα using RNAi mimicked the phenotype associated with loss of the bioactive peptides PETH and ETH (pre-ecdysis and ecdysis-triggering hormones, respectively) from Inka cells in *Drosophila*, suggesting that the Sytα contribute to neuropeptide release from neuroendocrine cells^93^. In addition, Seibert et al. recently reported that Syt9 may be required for the release of substance P from dense-core vesicles (DCVs) in striatal neurons in verterbrates^83^. Notably, both Syt1 and Syt7 are reported to play a role in DCV fusion in hippocampal neurons^94^, suggesting they may have multiple roles in regulating neurosecretion. We found that Syt1 mediates the fast ACh release and Syt7/Sytα mediates the slow sNPF release. Similarly, it has been shown in mouse neurons that Syt1 and Syt7 mediate the synchronous (fast) and asynchronous (slow) glutamate release, respectively^80^. Interestingly, Syt4, which does not contain a Ca^2+^-binding site, has been shown to negatively regulate the release of brain-derived neurotrophic factor (BDNF)^95^, while Syt10, which does contain a Ca^2+^-binding site, positively regulates the release of insulin-like growth factor 1 (IGF-1) from DCVs in neurons^96^. Together with our findings, these results support the notion that Syts have divergent roles and are involved in controlling distinct secretion pathways in neurons, depending on the specific cell type. Moreover, our results provide direct evidence that two Syt isoforms mediate neuropeptide release in *Drosophila*.

Why two Syt isoforms are required for the release of sNPF in the same neuron remains unclear. However, one possible explanation is that these two Syt isoforms function in the same secretory pathway. In this respect, it is interesting to note that previous studies suggested that Sytα may be localized to LDCVs^93^, while Syt7 may localized primarily to the peri-active zone^82^, and the results of Syt7 and Sytα double knock out also supports this conclusion.

In conclusion, we show that sNPF1.0 sensor is a robust tool for monitoring sNPF release *in vivo* with high specificity and spatiotemporal resolution. Our findings regarding the dynamics and molecular regulation of sNPF release provide valuable insights into the complex mechanisms by which neuropeptides and small molecule neurotransmitters are released from the same type of neurons.

## ACKNOWLEDGMENTS

We thank Yi Rao for providing access to the two-photon microscope. We thank the imaging core facility of State Key Laboratory of Membrane Biology at Peking University (Ye Liang), and Olympus/Evident China Life Science (Shaoling Qi, Wei Cao, Haitao Zhang, Dezhi Zhang, Linliang Yin, and Donghua Wu). We thank X. Lei at PKU-CLS and the National Center for Protein Sciences at Peking University for support and assistance with the Opera Phenix high-content screening system. We thank the Core Facility of Drosophila Resource and Technology of CAS Center for Excellence in Molecular Cell Science (Wei Wu). We thank members of the Li lab for helpful suggestions and comments on the manuscript. We thank Dr. Man jiang, Dr. Isabel Beets, Dr. Yufeng Pan, Dr. Yan Li, Dr. Xihuimin Dai and Dr. Edwin Levitan for valuable feedback of the manuscript.

This work was supported by grants from the Peking-Tsinghua Center for Life Sciences and the State Key Laboratory of Membrane Biology at Peking University School of Life Sciences, the National Natural Science Foundation of China (31925017 and 31871087 to Y.L.), the NIH BRAIN Initiative (NINDS U01NS120824 to Y.L.), the Feng Foundation of Biomedical Research, the Clement and Xinxin Foundation, and the New Cornerstone Science Foundation through the New Cornerstone Investigator Program and the XPLORER PRIZE (to Y.L.).

## AUTHOR CONTRIBUTIONS

Y.L. and X.X designed and supervised the project. X.X. performed and analyzed all experiments. Both authors analyzed and discussed the results. X.X. and Y.L. wrote the manuscript.

## STAR Methods

### EXPERIMENTAL MODEL AND SUBJECT DETAILS MATERIALS

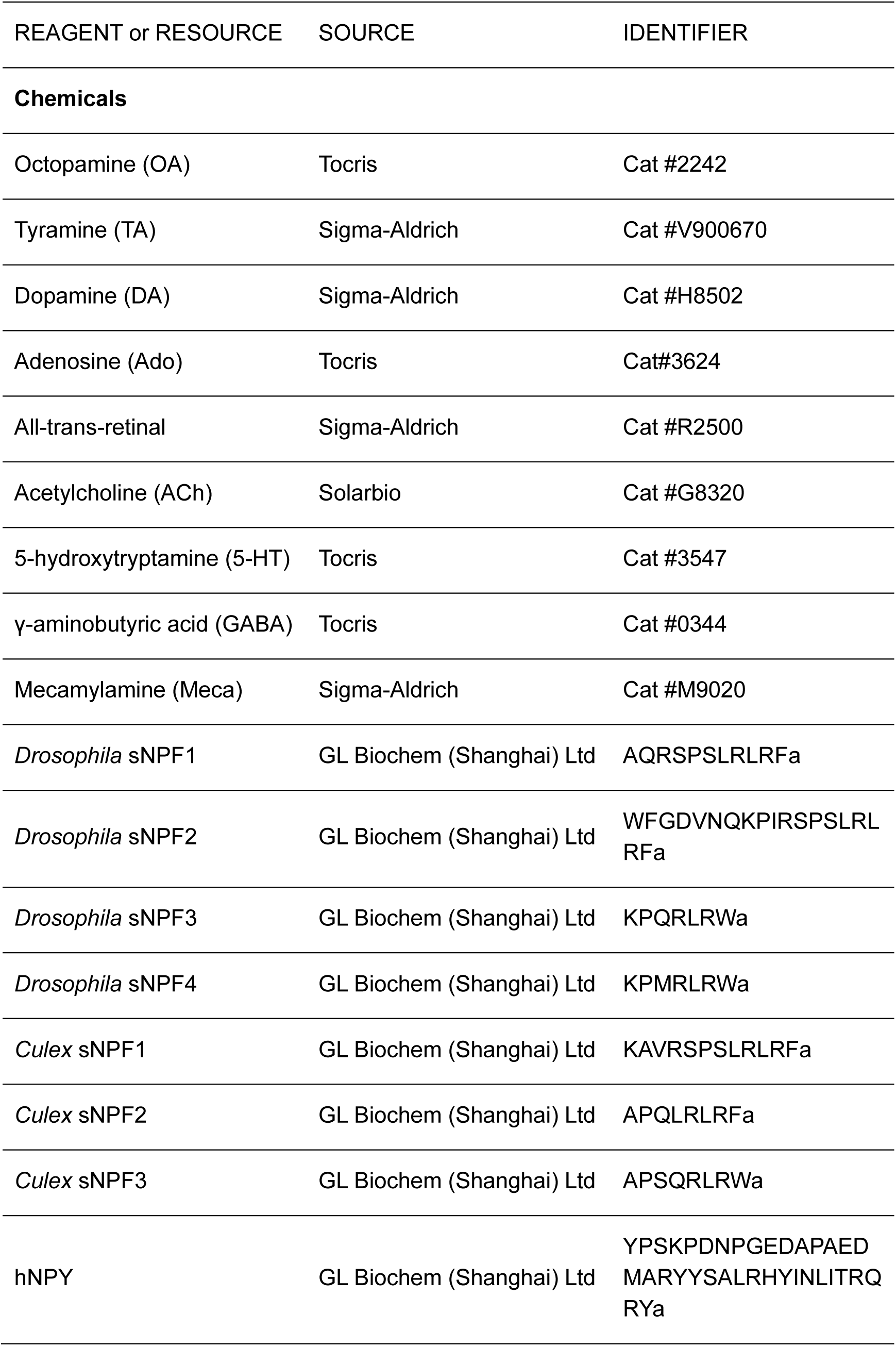
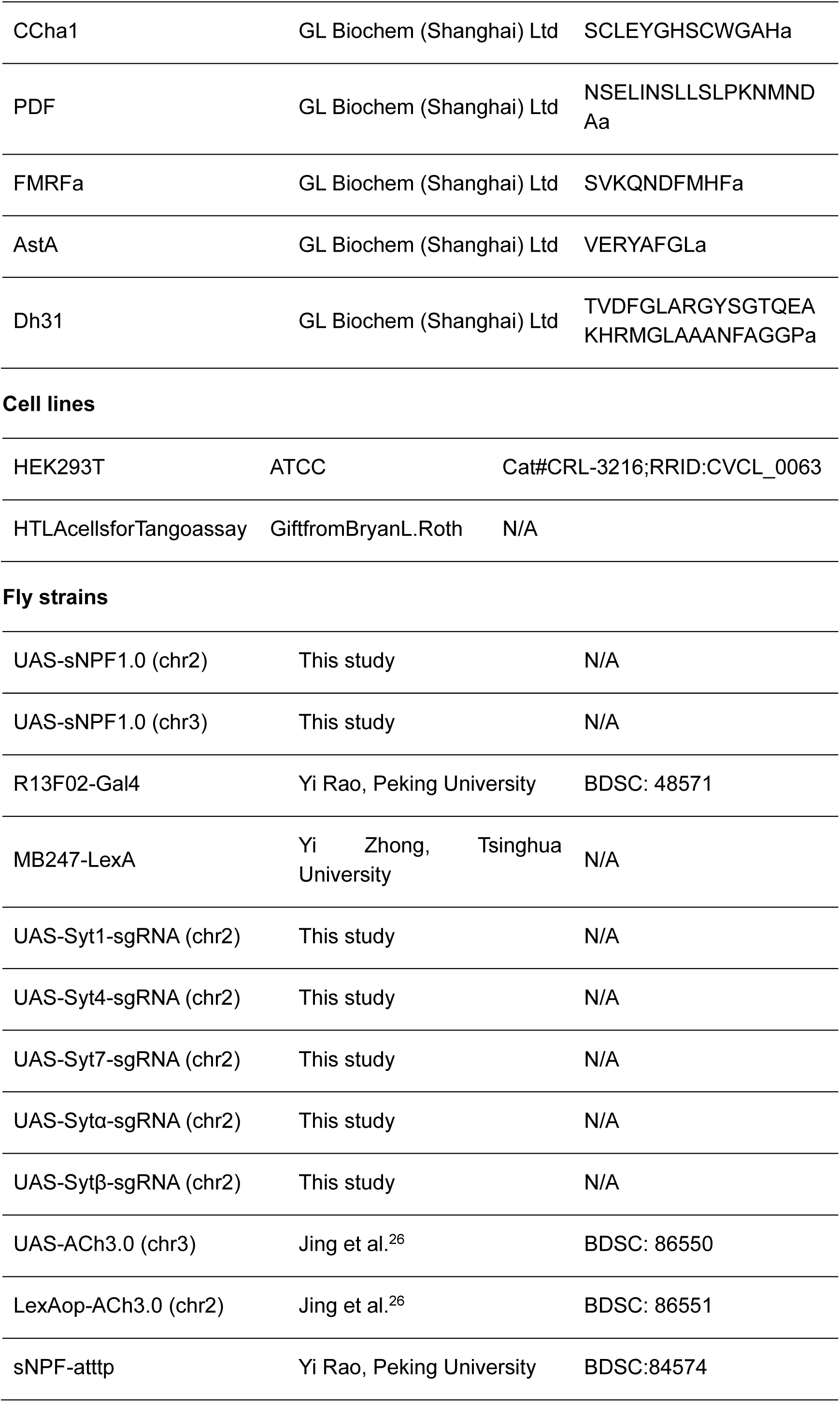
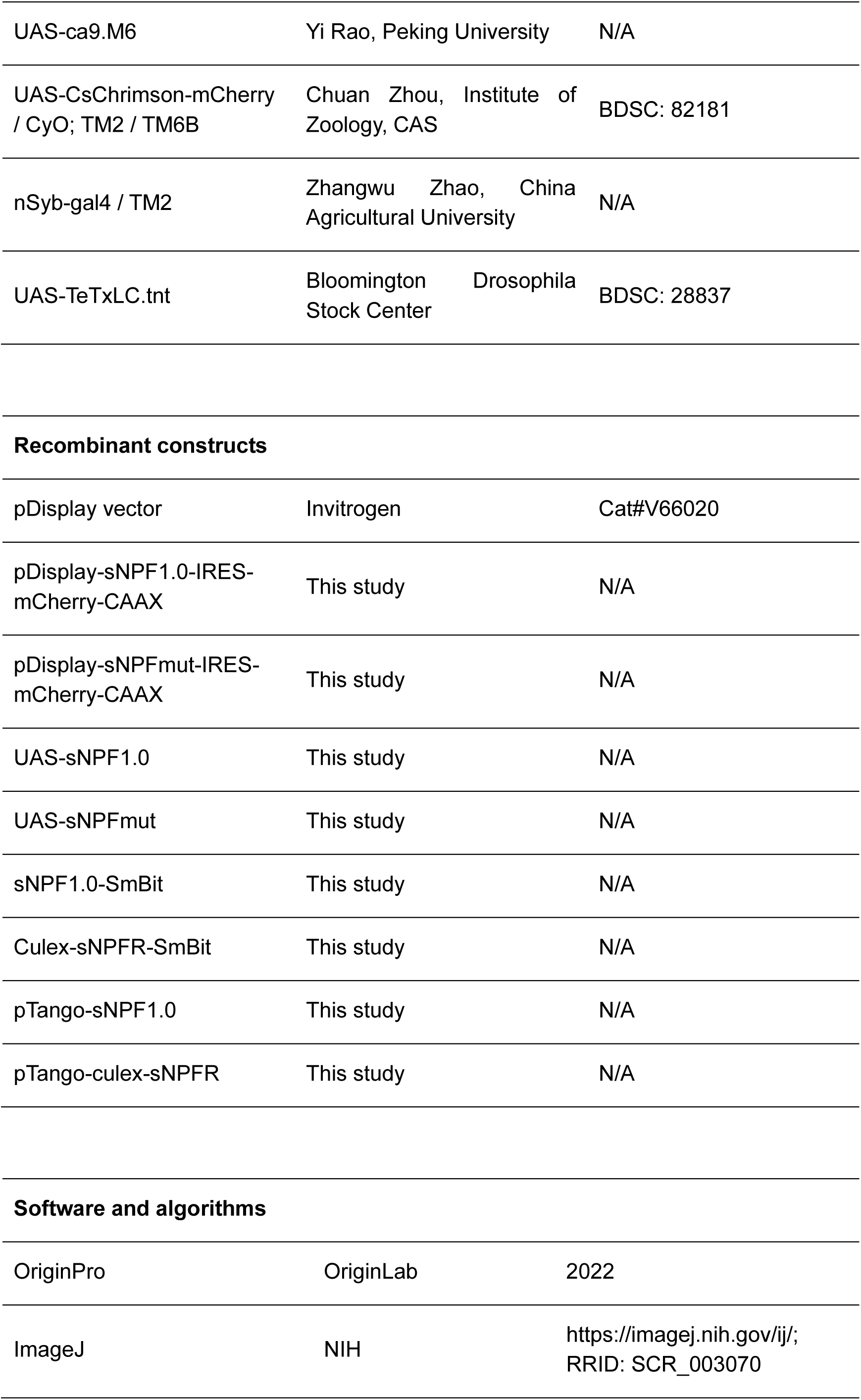
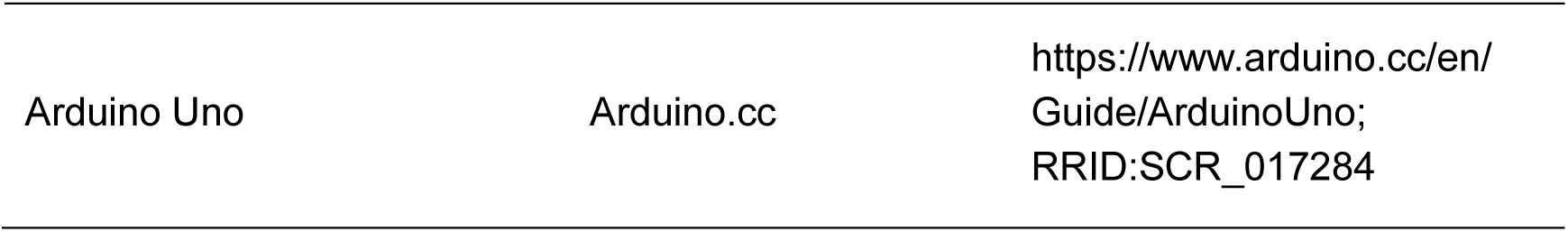

### METHOD DETAILS

#### Molecular cloning

The plasmids used in this study were generated using the Gibson assembly method. DNA inserts were generated by PCR amplification using primers (RuiBiotech) with ∼25-bp overlap, and all sequences were verified using Sanger sequencing (RuiBiotech). All cDNAs encoding the candidate GRAB_sNPF_ sensors were cloned into the pDisplay vector (Invitrogen) with an upstream IgK leader sequence and a downstream IRES-mCherry-CAAX cassette (to visualize localization to the cell membrane). For screening replacement sites, cDNAs encoding the various sNPF receptors were generated (Shang Genegay Biotech), and the third intracellular loop (ICL3) of each sNPF receptor was replaced with the corresponding ICL3 in GRAB_NE1m_. For optimizing the sNPF sensor, we screened the replaced sites in the *Culex* sNPF receptor, the amino acid composition between the *Culex* sNPF receptor and the ICL3 of GRAB_NE1m_, and cpEGFP. Site-directed mutagenesis was performed using primers containing randomized NNB codons (48 codons in total, encoding all 20 amino acids) or defined codons at the target sites.

#### Cell lines

HEK293T cells were acquired from ATCC and verified by microscopic examination of their morphology and growth curve. An HTLA cell line stably expressing a tTA-dependent luciferase reporter and the β-arrestin2-TEV fusion gene used in the Tango assay was a generous gift from Bryan L. Roth (University of North Carolina Chapel Hill). The cells were cultured in DMEM (Biological Industries) supplemented with 10% (v/v) fetal bovine serum (FBS, Gibco) and 1% penicillin-streptomycin (Gibco) at 37°C in humidified air containing 5% CO_2_.

#### Fly strain generation and animal husbandry

In this study, we generated UAS-sNPF1.0 (attp40, UAS-sNPF1.0/CyO), UAS-sNPF1.0 (vk00005, UAS-sNPF1.0/TM2), and UAS-sNPFmut (attp40, UAS-sNPFmut/CyO) vectors using Gibson assembly to integrate the coding sequence of sNPF1.0 into the pJFRC28 (Addgene plasmid 36431) or modified pJFRC28 vector.

The UAS-Syt1-sgRNA, UAS-Syt4-sgRNA, UAS-Syt7-sgRNA, UAS-Sytα-sgRNA, and UAS-Sytβ-sgRNA constructs were designed by inserting three guide RNAs (sgRNAs) into the pMsgNull vector based on pACU2 (Addgene #31223)^97^ (From Dr. Yi Rao lab at Peking University), with rice transfer RNA (tRNA) used to separate the various sgRNAs. The resulting vectors were then injected into embryos and integrated into attp40 or vk00005 via phiC31 by the Core Facility of Drosophila Resource and Technology, Shanghai Institute of Biochemistry and Cell Biology, Chinese Academy of Sciences.

The flies were raised on standard corn meal–yeast medium at 25°C in 50% relative humidity under a 12h/12h light/dark cycle. For optogenetics, after eclosion, the flies were transferred to corn meal containing 400 μM all-*trans*-retinal and raised in the dark for 1-3 days before performing functional imaging experiments.

#### sgRNA sequences

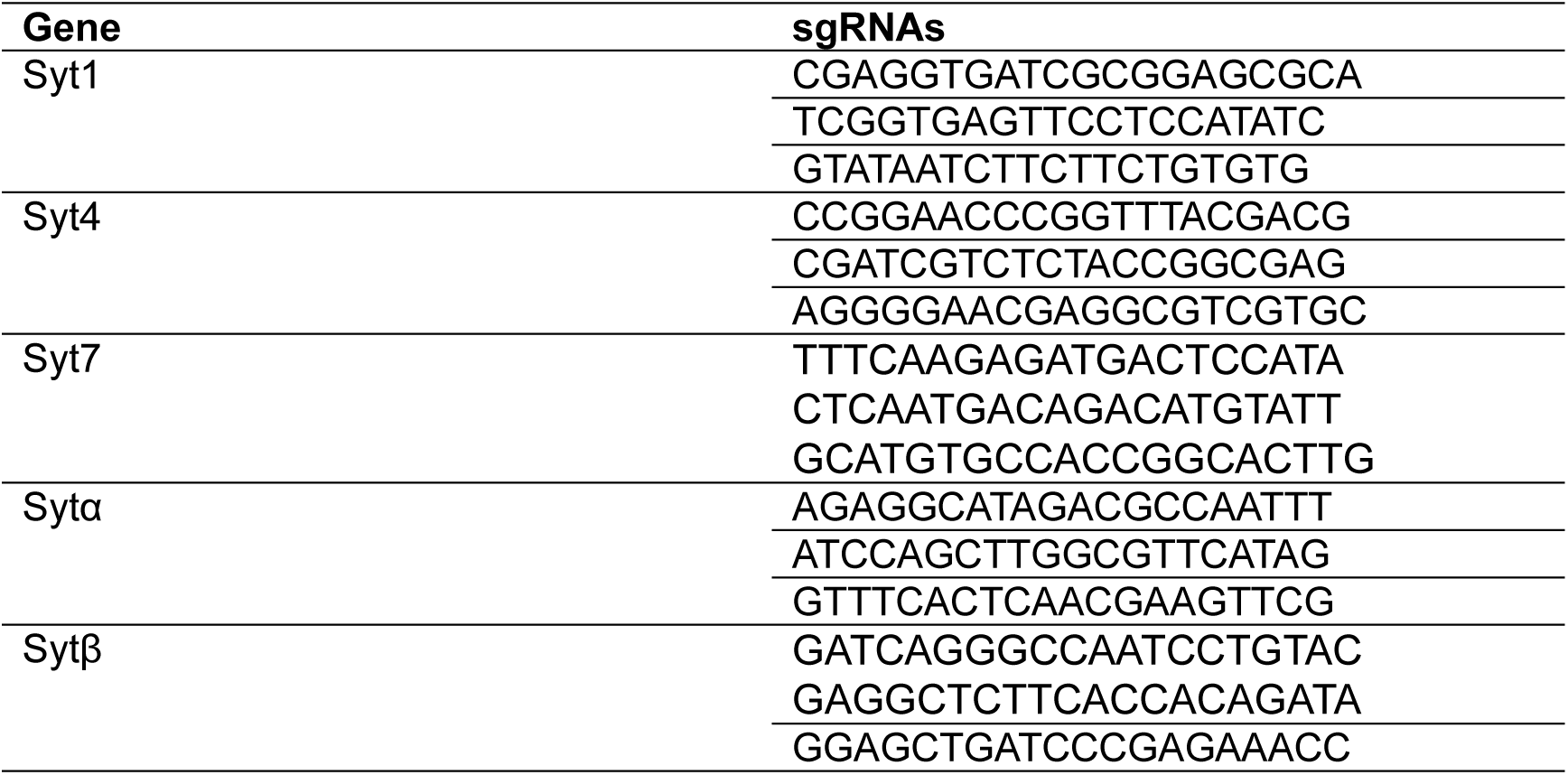

#### Fly genotypes used in each figure

| Figure | Genotype |
| --- | --- |
| Fig. 1 |  |
| 1I-K | UAS-sNPF1.0 / CyO; R13F02-Gal4 / TM2 |
| Fig. 2 |  |
| 2A-C | UAS-sNPF1.0 / CyO; nsyb-Gal4 / TM2 |
| 2A-C | UAS-sNPF1.0 / sNPF-attp; nsyb-Gal4 / sNPF-attp |
| 2E-K | UAS-sNPF1.0 / UAS-CsChrimson-mCherry; R13F02-Gal4 / TM2 |
| Fig. 3 |  |
| 3B, D-E | UAS-sNPF1.0 / UAS-CsChrimson-mCherry; R13F02-Gal4 / TM2 |
| 3C, D-E | LexAop-ACh3.0 / UAS-CsChrimson-mCherry; R13F02-Gal4, Mb247-LexA / TM2 |
| Fig. 4 |  |
| 4B-C, F-G, | UAS-sNPF1.0 / UAS-CsChrimson-mCherry; R13F02-Gal4 / TM2 |
| 4D-G, | LexAop-ACh3.0 / UAS-CsChrimson-mCherry; R13F02-Gal4, Mb247-LexA / TM2 |
| 4H | UAS-sNPF1.0 / UAS-CsChrimson-mCherry; R13F02-Gal4 / TM2 |
| 4I | UAS-ACh3.0 / UAS-CsChrimson-mCherry; R13F02-Gal4 / TM2 |
| Fig. 5 |  |
| 5C, E, H-J | UAS-sNPF1.0 / UAS-CsChrimson-mCherry; R13F02-Gal4 / TM2 |
| 5D, F, K-M | UAS-ACh3.0 / UAS-CsChrimson-mCherry; R13F02-Gal4 / TM2 |
| Fig. 6 |  |
| 6A | LexAop-ACh3.0 / UAS-CsChrimson-mCherry; R13F02-Gal4, Mb247-LexA / TM2 |
| 6A | LexAop-ACh3.0 / UAS-CsChrimson-mCherry; R13F02-Gal4, Mb247-LexA / UAS-TeTxLC.tnt |
| 6B | UAS-sNPF1.0 / CyO; R13F02-Gal4 / TM2 |
| 6B | UAS-sNPF1.0 / UAS-TeTxLC.tnt; R13F02-Gal4 / + |
| 6C, E | UAS-cas9.M6, UAS-CsChrimson-mCherry / CyO; R13F02-Gal4, UAS-sNPF1.0 / TM2 |
| 6C, E | UAS-cas9.M6, UAS-CsChrimson-mCherry / UAS-Syt1-sgRNA; R13F02-Gal4, UAS-sNPF1.0 / + |
| 6C, E | UAS-cas9.M6, UAS-CsChrimson-mCherry / UAS-Syt4-sgRNA; R13F02-Gal4, UAS-sNPF1.0 / + |
| 6C, E | UAS-cas9.M6, UAS-CsChrimson-mCherry / UAS-Syt7-sgRNA; R13F02-Gal4, UAS-sNPF1.0 / + |
| 6C, E | UAS-cas9.M6, UAS-CsChrimson-mCherry / UAS-Sytα-sgRNA; R13F02-Gal4, UAS-sNPF1.0 / + |
| 6C, E | UAS-cas9.M6, UAS-CsChrimson-mCherry / UAS-Sytβ-sgRNA; R13F02-Gal4, UAS-sNPF1.0 / + |
| 6D, F | UAS-cas9.M6, UAS-CsChrimson-mCherry / CyO; R13F02-Gal4, UAS-ACh3.0 / TM2 |
| 6D, F | UAS-cas9.M6, UAS-CsChrimson-mCherry / UAS-Syt1-sgRNA; R13F02-Gal4, UAS-ACh3.0 / + |
| 6D, F | UAS-cas9.M6, UAS-CsChrimson-mCherry / UAS-Syt4-sgRNA; R13F02-Gal4, UAS-ACh3.0 / + |
| 6D, F | UAS-cas9.M6, UAS-CsChrimson-mCherry / UAS-Syt7-sgRNA; R13F02-Gal4, UAS-ACh3.0 / + |
| 6D, F | UAS-cas9.M6, UAS-CsChrimson-mCherry / UAS-Sytα-sgRNA; R13F02-Gal4, UAS-ACh3.0 / + |
| 6D, F | UAS-cas9.M6, UAS-CsChrimson-mCherry / UAS-Sytβ-sgRNA;<br>R13F02-Gal4, UAS-ACh3.0 / + |
| Fig. S3 |  |
| 3A-G | UAS-sNPF1.0 / UAS-CsChrimson-mCherry; R13F02-Gal4 / TM2 |
| Fig. S4 |  |
| 4A, B | UAS-sNPF1.0 / UAS-CsChrimson-mCherry; R13F02-Gal4 / TM2 |
| 4C, D | LexAop-ACh3.0 / UAS-CsChrimson-mCherry; R13F02-Gal4, Mb247-LexA / TM2 |
| Fig. S5 |  |
| 5A, C-D | UAS-sNPF1.0 / UAS-CsChrimson-mCherry; R13F02-Gal4 / TM2 |
| 5B, C-D | LexAop-ACh3.0 / UAS-CsChrimson-mCherry; R13F02-Gal4, Mb247-LexA / TM2 |
| Fig. S6 |  |
| 6A, B | UAS-cas9.M6, UAS-CsChrimson-mCherry / UAS-Sytα-sgRNA;<br>R13F02-Gal4, UAS-sNPF1.0 / + |
| 6A, B | UAS-cas9.M6, UAS-CsChrimson-mCherry / UAS-Syt7-sgRNA;<br>R13F02-Gal4, UAS-sNPF1.0 / + |
| 6A, B | UAS-cas9.M6, UAS-CsChrimson-mCherry / UAS-Sytα-sgRNA;<br>R13F02-Gal4, UAS-sNPF1.0 / UAS-Syt7-sgRNA |

#### Fluorescence imaging of HEK 293T cells

Cells were imaged using an inverted Ti-E A1 confocal microscope (Nikon) or an Opera Phenix high-content screening system (PerkinElmer). The confocal microscope was equipped with a 10x/0.45 NA (numerical aperture) objective, a 20x/0.75 NA objective, a 40x/1.35 NA oil-immersion objective, a 488-nm laser, and a 561-nm laser; the GFP signal was collected using a 525/50-nm emission filter combined with the 488-nm laser, and the RFP signal was collected using a 595/50-nm emission filter combined with the 561-nm laser. The Opera Phenix system was equipped with 20x/0.4 NA objective, a 40x/1.1 NA water-immersion objective, a 488-nm laser, and a 561-nm laser; the GFP and RFP signals were collected using a 525/50-nm and 600/30-nm emission filter, respectively. The fluorescence signal produced by the green fluorescent GRAB_sNPF_ sensors was calibrated using the GFP/RFP ratio.

HEK293T cells were plated on either 12-mm glass coverslips in 24-well plates or 96-well plates and grown to ∼70% confluence for transfection with PEI (1 μg plasmid and 3 μg PEI per well in 24-well plates or 300 ng plasmids and 900 ng PEI per well in 96-well plates); the medium was replaced after 4–6 hours, and the cells were used for imaging 24–48 h after transfection. To measure the kinetics of the GRAB_sNPF_ sensor, the confocal high-speed line scanning mode (1024 Hz) was used to measure the fluorescence signal change when the cells were locally puffed with sNPF via a glass pipette positioned in close proximity to the cells, the increased trace in fluorescence was fitted with a single-exponential function.

#### Tango assay

HTLA cells were cultured in 6-well plates; at ∼70% cell density, the cells were transfected with either wild-type *Culex* sNPFR or the sNPF1.0 sensor. Twenty-four hours after transfection, the cells were transferred to a 96-well white clear flat-bottom plate, and various concentrations of sNPF (ranging from 0.1 nM to 5 μM) were added to the cells; each concentration was applied in triplicate. The cells were then incubated for ∼16 hours, and the bioluminescent signal was measured. To measure the bioluminescent signal, the culture medium was removed, and 40 μl of Bright-Glo substrate (Promega) was added to each well. The plate was then incubated at room temperature in the dark for 10 minutes, and bioluminescence was measured using a Victor X5 microplate reader (PerkinElmer). Non-transfected cells were used a negative control.

#### Luciferase complementation assay

The luciferase complementation assay was performed as previously described^98^. In brief, 24–48 h after transfection, the cells were washed with PBS, dissociated using a cell scraper, resuspended in PBS, transferred to opaque 96-well plates containing 5 μM furimazine (NanoLuc Luciferase Assay, Promega), and bathed in sNPF at various concentrations (ranging from 0.1 nM to 5 μM). After incubation for 10 minutes in the dark, luminescence was measured using a Victor X5 microplate reader (PerkinElmer).

#### Spectra measurements

For one-photon spectra, HEK293T cells were transfected with CMV promoter‒driven sNPF1.0 plasmids; after 24 h, the cells were harvested and transferred to a 384-well plate in the absence or presence of 1 µM sNPF. Excitation and emission spectra were measured at 5-nm increments with a 20-nm bandwidth using a Safire2 multi-mode plate reader (Tecan). For background subtraction, non-transfected cells were prepared and measured using the same protocol.

For two-photon spectra, cells were transfected with sNPF1.0 and treated as described above. Excitation and emission spectra were measured from 700 nm to 1020 nm at 10-nm increments using an FV1000 two-photon microscope (Olympus) equipped with a Spectra-Physics Mai Tai Ti:Sapphire laser. Non-transfected cells were used to subtract the background signal.

#### Two-photon imaging of flies

Fluorescence imaging in flies was performed using an FV1000 two-photon microscope (Olympus) equipped with a Spectra-Physics Mai Tai Ti:Sapphire laser. A 920-nm excitation laser was used for one-color imaging of sNPF1.0 and sNPFmut, and a 950-nm excitation laser was used for two-color imaging of sNPF1.0 and mCherry. For detection, a 495-540-nm filter was used for the green channel, and a 575-630-nm filter was used for red channel. Adult female flies were used for imaging within 1 week after eclosion. To prepare the fly for imaging, adhesive tape was affixed to the head and wings. The tape above the head was excised, and the chitin head-shell, air sacs, and fat bodies were carefully removed to expose the central brain. The brain was bathed continuously in an adult hemolymph-like solution composed of (in mM): 108 NaCl, 5 KCl, 5 HEPES, 5 trehalose, 5 sucrose, 26 NaHCO_3_, 1 NaH_2_PO_4_, 2 CaCl_2_, and 1-2 MgCl_2_. For single-photon optogenetic stimulation, a 635-nm laser (Changchun Liangli Photo Electricity Co., Ltd.) was used, and 18 mW/cm^2^ light pulses were delivered to the brain via an optic fiber. For the perfusion experiments, a small section of the blood-brain-barrier was carefully removed with tweezers before applying the indicated compounds or solutions.

#### Quantification and statistical analysis

##### Imaging experiments

Images were processed using ImageJ software (National Institutes of Health). The change in fluorescence (ΔF/F_0_) was calculated using the formula [(F-F_0_)/F_0_], where F_0_ represents the baseline fluorescence. The signal-to-noise ratio (SNR) was calculated by dividing the peak response by the standard deviation of the baseline fluorescence. The area under the curve was determined using the integral of the change in fluorescence (∫ΔF/F_0_).

##### Statistical analysis

Origin 2019 (OriginLab) was used to perform the statistical analyses. Unless otherwise specified, all summary data are presented as the mean ± sem. The paired or unpaired Student’s *t*-test was used to compare two groups, and a one-way analysis of variance (ANOVA) was used to compare more than two groups. All statistical tests were two-tailed, and differences were considered statistically significant at *P* < 0.05.

##### Code availability

The custom-written R, Arduino, and ImageJ programs will be provided upon request.

---

**Fig. S1.**
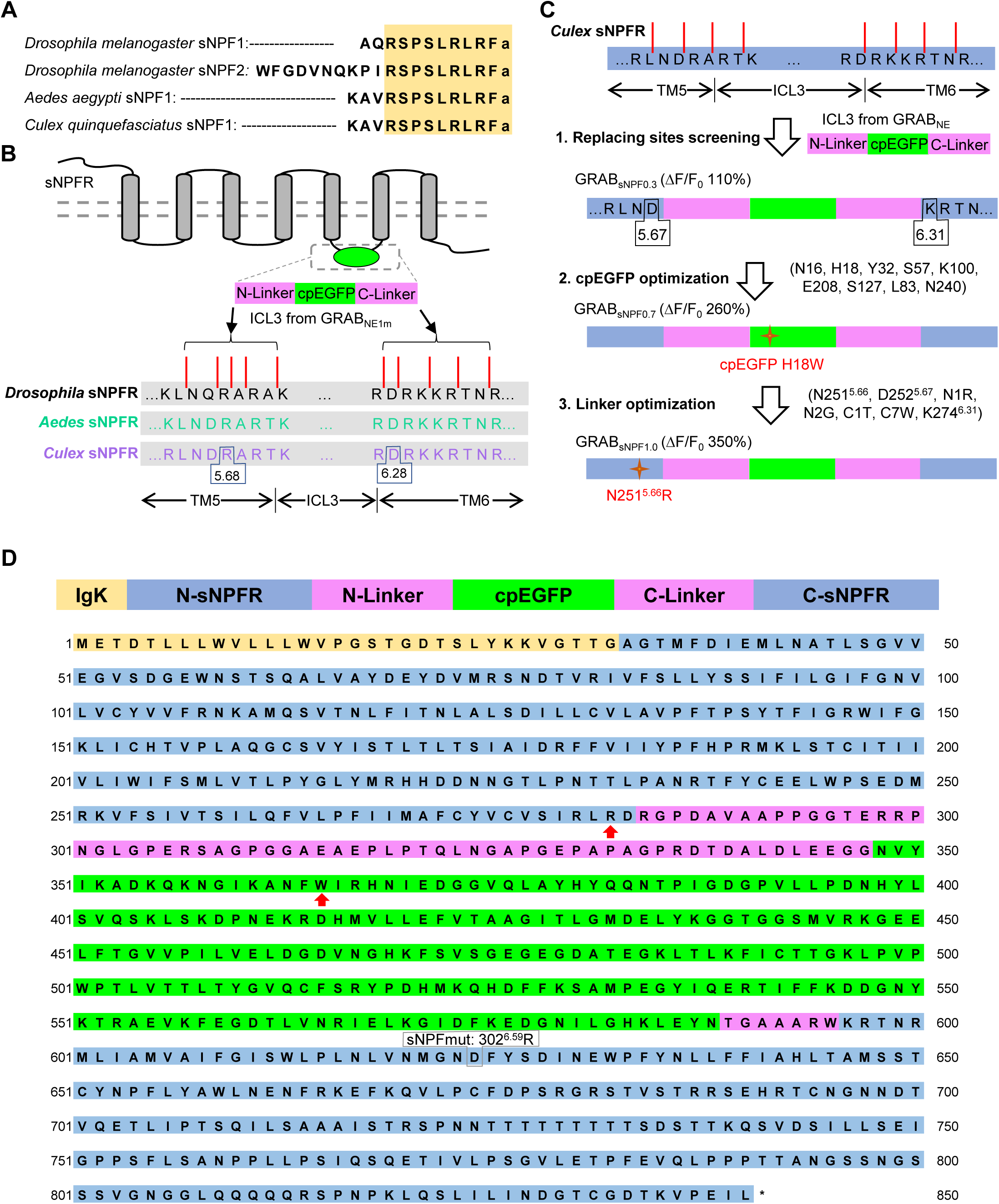
Strategy for optimizing and screening GRAB_sNPF_ sensors. (A) Alignment of the amino acid sequences of *Drosophila melanogaster*, *Aedes aegypti*, and *Culex quinquefasciatus* sNPF1. (B) Schematic diagram depicting the strategy for replacing the indicated sites for screening using the indicated sNPF receptors. (C) Flowchart depicting the steps used for developing and optimizing the sNPF1.0 GRAB sensor. (D) Top, schematic diagram depicting the structural features of the GRAB_sNPF1.0_ sensor, showing the IgK leader sequence, the N-terminal and C-terminal sNPFR-derived sequences, and cpEGFP with flanking linker domains. Bottom, amino acid sequence of the sNPF1.0 sensor. Note that the numbering system used in this figure corresponds to the start of the IgK leader sequence. Red arrows indicate mutated amino acids, and the position of the point mutation to generate the ligand-insensitive sensor, sNPFmut, is indicated.

**Fig. S2.**
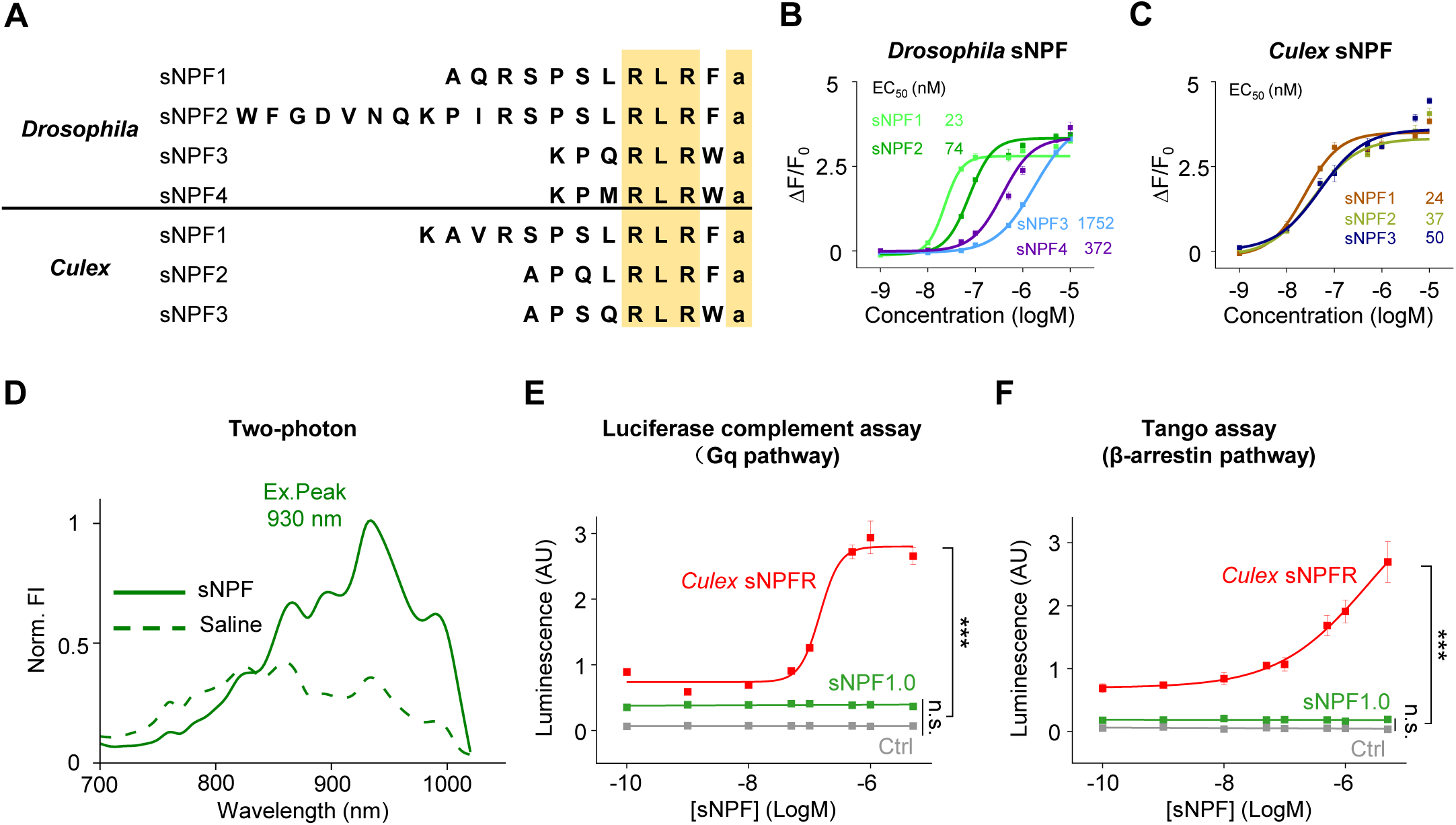
Characterization of GRAB_sNPF1.0_ sensor in HEK293T cells. (A) Alignment of the amino acid sequences of the major sNPF isoforms in *Drosophila* and *Culex*. (B and C) Dose‒response curves for sNPF1.0 expressed in HEK293T cells in response to increasing concentrations of the indicated *Drosophila* (B) and *Culex* (C) sNPF isoforms, with the corresponding EC_50_ values shown; n = 3 wells with 200‒400 cells per well. (A) (D) Two-photon excitation spectra of sNPF1.0 measured in the absence and presence of sNPF. (B) (E) Summary of relative dose-dependent downstream G protein coupling in control HEK293T cells and in cells expressing either wild-type *Culex* sNPFR or sNPF1.0, measured using the luciferase complementation mini-G protein assay; n = 3 wells per group, 200–500 cells per well. (C) (F) Summary of relative dose-dependent downstream β-arrestin coupling in control HEK293T cells and in cells expressing either wild-type *Culex* sNPFR or sNPF1.0, measured using the Tango assay; n = 3 wells per group, 200–500 cells per well. Data are shown as mean ± s.e.m. \*\*\**P* < 0.001 and n.s., not significant.

**Fig. S3.**
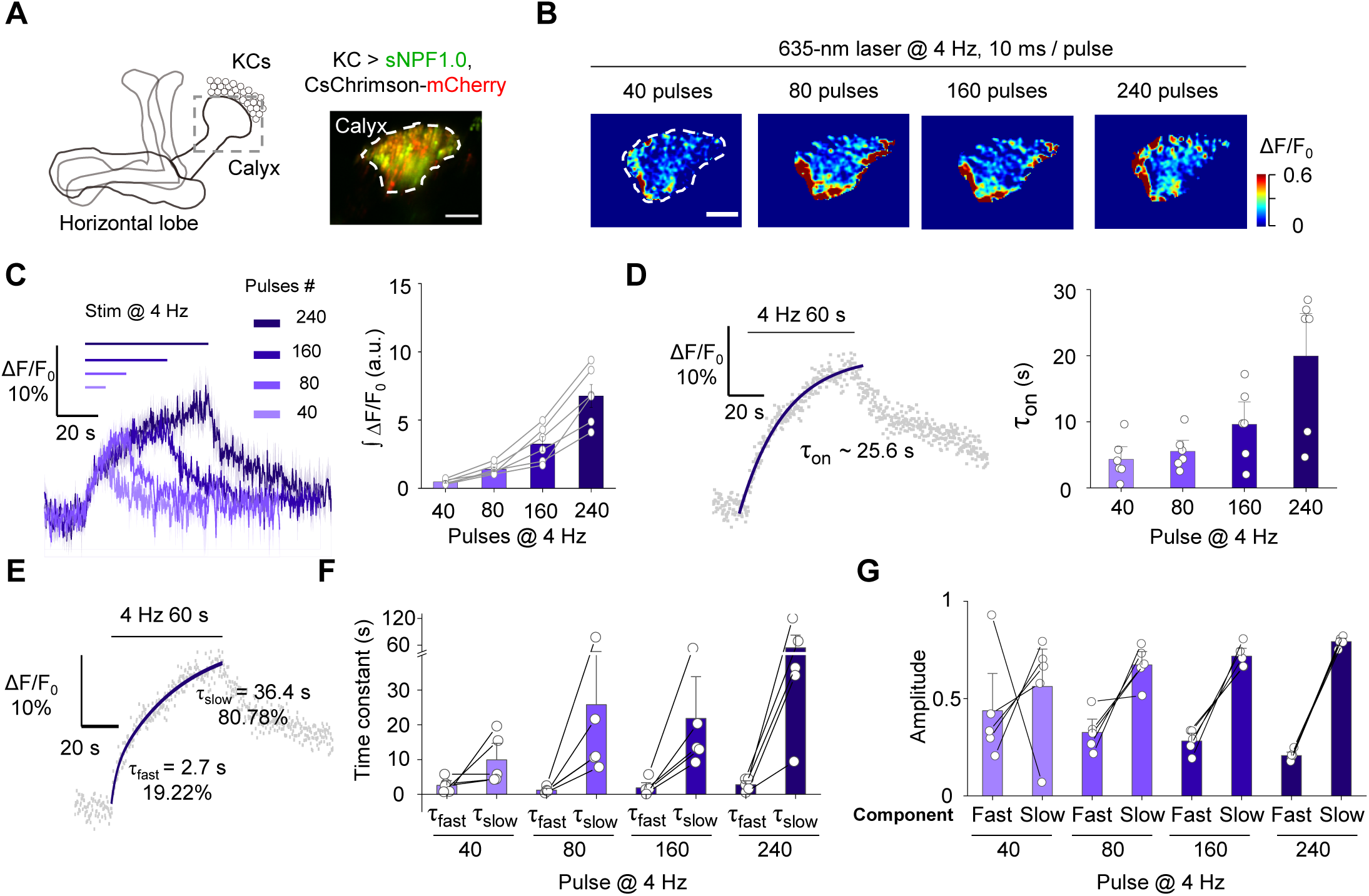
The GRAB_sNPF_ sensor can report sNPF release in the dendritic region in the *Drosophila* MB. (A) Schematic diagram depicting the experimental setup (left) and an example fluorescence image (right) of sNPF1.0 and CsChrimson-mCherry in the calyx region. (B and C) Representative pseudocolor images (B), traces (C, left), and summary (C, right) of the change in sNPF1.0 fluorescence in response to the indicated number of 635-nm laser pulses applied at 4 Hz; n = 6 flies. Scale bars, 25 μm. (A) (D) Left, sNPF1.0 fluorescence was measured before, during, and after 240 pulses of 635-nm light, and the rise phase was fitted with a single-exponential function. Right, summary of the rise time constant; n = 6 flies. (B) (E) sNPF1.0 fluorescence was measured before, during, and after 240 pulses of 635-nm light, and the rise phase was fitted with a double-exponential function. (F and G) Summary of the rise time constants (F) and amplitudes (G); n = 6 flies. Data are shown as mean ± s.e.m. in c, with the error bars or shaded regions indicating the s.e.m.

**Fig. S4.**
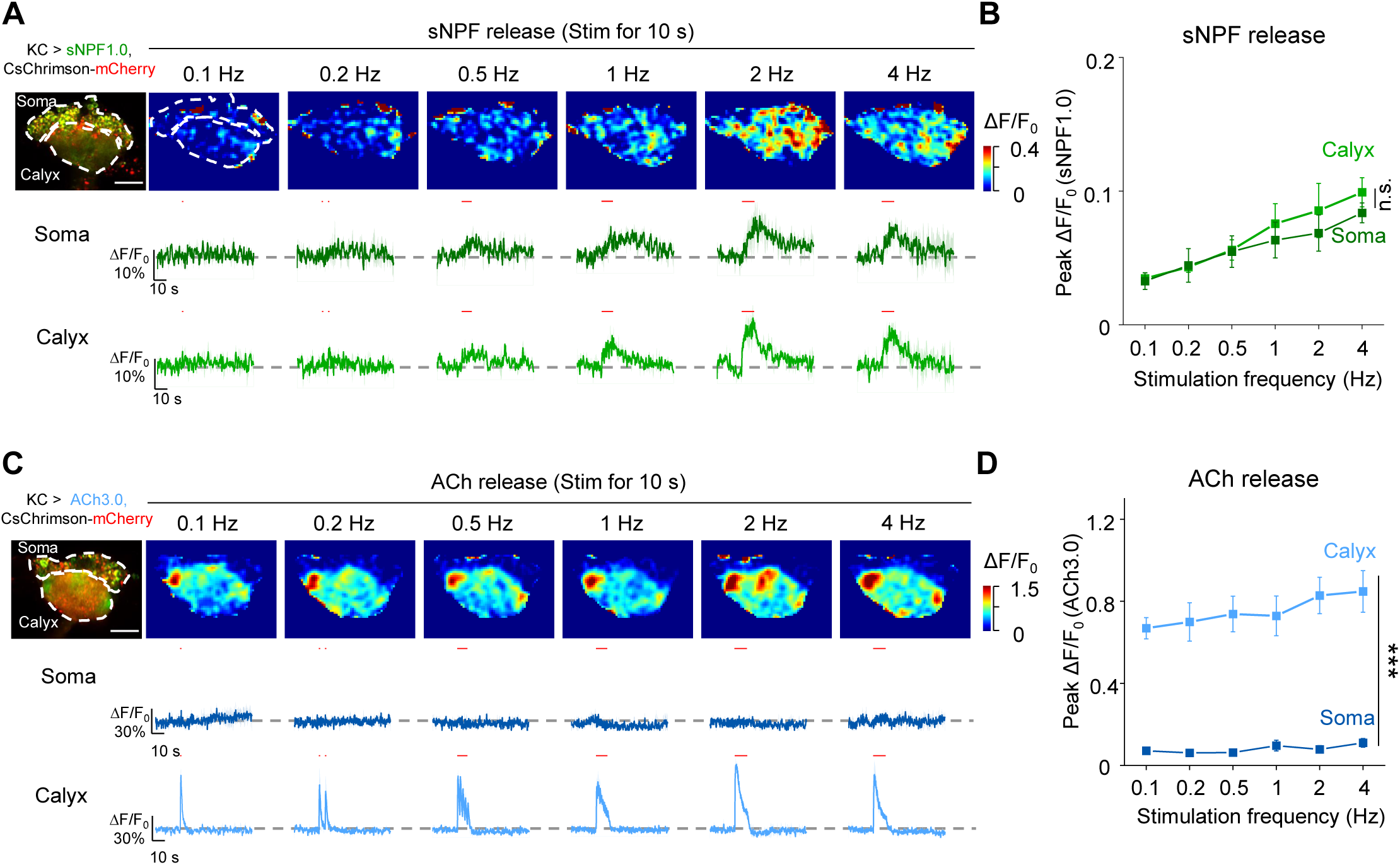
GRAB sensors reveal spatially distinct patterns of sNPF and ACh release from KCs. (A and B) Representative fluorescence image (A, top left), pseudocolor images (A, top right), traces (A, bottom right), and summary (B) of the change in sNPF1.0 fluorescence measured in the soma and calyx in response to the indicated frequencies of 635-nm laser pulses applied for 10 s; n = 5 flies per group. Scale bars, 25 μm. The 100 μM nAChR antagonist mecamylamine (Meca) was present throughout these experiments. (C and D) Representative fluorescence image (C, top left), pseudocolor images (C, top right), traces (C, bottom right), and summary (D) of the change in ACh3.0 fluorescence measured in the soma and calyx in response to the indicated frequencies of 635-nm laser pulses applied for 10 s; n = 6-7 flies. Data are shown as mean ± s.e.m. in a and c, with the error bars or shaded regions indicating the s.e.m. \*\*\**P* < 0.001 and n.s., not significant.

**Fig. S5.**
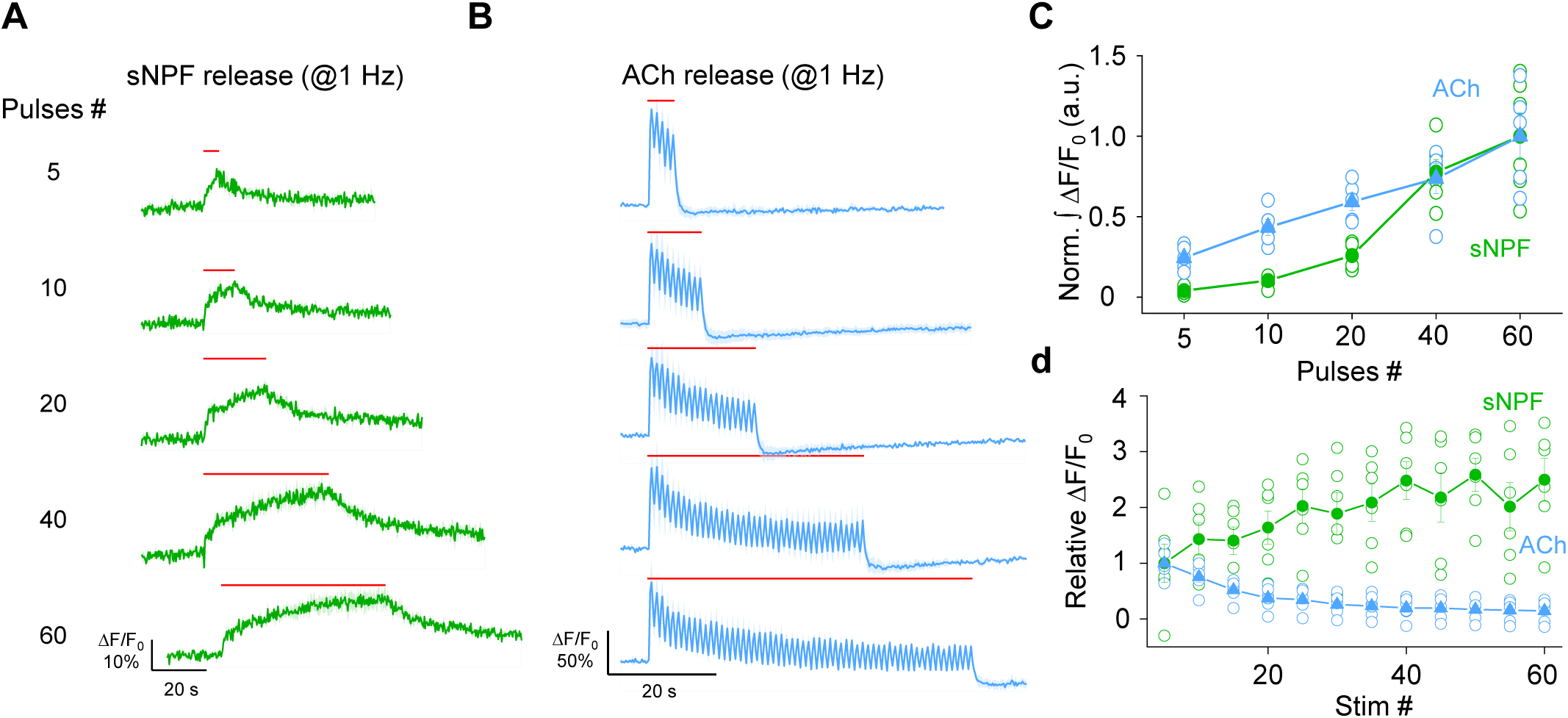
GRAB sensors reveal differences in activity-dependent dynamics between sNPF and ACh release from KCs. (A and B) Representative traces of the change in sNPF1.0 (A) and ACh3.0 (B) fluorescence in response to the indicated number of light pulses applied at 1 Hz. The 100 μM nAChR antagonist mecamylamine (Meca) was present throughout these experiments. (C and D) Summary of normalized integrated ΔF/F_0_ (C) and relative ΔF/F_0_ (D) for sNPF1.0 and ACh3.0 measured in response to the indicated number of light pulses applied at 1 Hz. Data are shown as mean ± s.e.m. in a and b, with the error bars or shaded regions indicating the s.e.m.

**Fig. S6.**
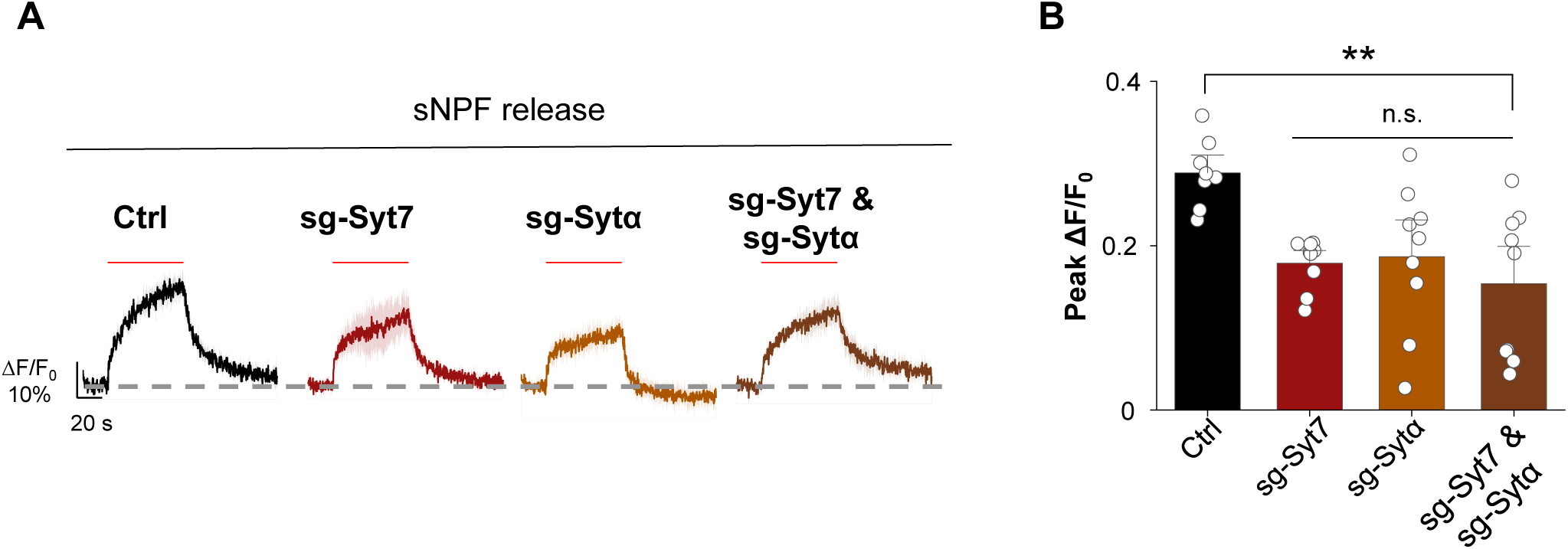
Knocking out both Syt7 and Sytα shows similar sNPF release with knocking out each synaptotagmin isoform individually. (A) Representative traces of the change in sNPF1.0 fluorescence in response to 240 light pulses applied at 4 Hz (red horizontal lines) in control flies (Ctrl), Syt7 knockout flies, Sytα knockout flies, and Syt7/Sytα double knock out flies. (B) Summary of the peak change in sNPF1.0 fluorescence measured in the indicated flies. Data are shown as mean ± s.e.m. in a, with the error bars or shaded regions indicating the s.e.m. \*\**P* < 0.01 and n.s., not significant.

